# SREBP-dependent regulation of lipid homeostasis is required for progression and growth of pancreatic ductal adenocarcinoma

**DOI:** 10.1101/2024.02.04.578802

**Authors:** Chiaki T. Ishida, Stephanie L. Myers, Wei Shao, Meredith R. McGuire, Chune Liu, Casie S. Kubota, Theodore E. Ewachiw, Debaditya Mukhopadhyay, Suqi Ke, Hao Wang, Zeshaan A. Rasheed, Robert A. Anders, Peter J. Espenshade

## Abstract

Metabolic reprogramming is a necessary component of oncogenesis and cancer progression that solid tumors undergo when their growth outstrips local nutrient supply. The supply of lipids such as cholesterol and fatty acids is required for continued tumor cell proliferation, and oncogenic mutations stimulate de novo lipogenesis to support tumor growth. Sterol regulatory element-binding protein (SREBP) transcription factors control cellular lipid homeostasis by activating genes required for lipid synthesis and uptake. SREBPs have been implicated in the progression of multiple cancers, including brain, breast, colon, liver, and prostate. However, the role the SREBP pathway and its central regulator SREBP cleavage activating protein (SCAP) in pancreatic ductal adenocarcinoma (PDAC) has not been studied in detail. Here, we demonstrated that pancreas-specific knockout of *Scap* has no effect on mouse pancreas development or function, allowing for examination of the role for *Scap* in the murine KPC model of PDAC. Notably, heterozygous loss of *Scap* prolonged survival in KPC mice, and homozygous loss of *Scap* impaired PDAC tumor progression. Using subcutaneous and orthotopic xenograft models, we showed that S*CAP* is required for human PDAC tumor growth. Mechanistically, chemical or genetic inhibition of the SREBP pathway prevented PDAC cell growth under low serum conditions due to a lack of lipid supply. Highlighting the clinical importance of this pathway, the SREBP pathway is broadly required for cancer cell growth, SREBP target genes are upregulated in human PDAC tumors, and increased expression of SREBP targets genes is associated with poor survival in PDAC patients. Collectively, these results demonstrate that SCAP and the SREBP pathway activity are essential for PDAC cell and tumor growth *in vitro* and *in vivo*, identifying SCAP as a potential therapeutic target for PDAC.

**SIGNIFICANCE:** Our findings demonstrate that SREBP pathway activation is a critical part of the metabolic reprogramming that occurs in PDAC development and progression. Therefore, targeting the SREBP pathway has significant therapeutic potential.

## INTRODUCTION

Pancreatic ductal adenocarcinoma (PDAC) is a lethal disease with a 5-year overall survival rate of 12% and an increasing incidence (1). Poor survival can be attributed to late diagnosis and a standard of care that prolongs survival by less than one year (2). Despite recent progress in translational advances for PDAC treatment, new strategies are still desperately needed to significantly improve clinical outcomes (3).

Cancer cells undergo metabolic reprogramming in response to oncogenic mutations and the tumor microenvironment (4). Lipids such as cholesterol and fatty acids are required to support membrane synthesis, cell signaling, and energetics, and lipid metabolic reprogramming has been identified as an important step in tumorigenesis (5). Cancer cells maintain lipid supplies by upregulating *de novo* synthesis and/or lipid uptake from the environment (5). Two genes, *SREBF1* and *SREBF2*, code for three endoplasmic reticulum (ER) membrane-bound transcription factors, SREBP-1a, SREBP-1c, and SREBP-2 that regulate cellular lipid homeostasis. SREBP-1 primarily regulates fatty acid and triglyceride synthesis, and SREBP-2 controls cholesterol synthesis enzymes and uptake through the low-density lipoprotein receptor (LDLR) (6). Each SREBP isoform is negatively regulated by lipid supply through the function of the ER integral membrane protein SREBP cleavage activation protein (SCAP). SREBPs form a tight complex with SCAP, which is retained in the ER under conditions of high lipid supply (7). Under low lipid conditions, the SREBP-SCAP complex traffics to the Golgi, where SREBP is sequentially cleaved by the site-1 (S1P) and site-2 (S2P) proteases. The N-terminal transcription factor domain of SREBP (SREBP-N) is released from the membrane and enters the nucleus, where it activates transcription of lipid metabolic genes such as *low-density lipoprotein receptor (LDLR), stearoyl-CoA desaturase (SCD),* and *HMG-CoA reductase* (*HMGCR*) (6). Upregulation of lipid synthesis and uptake restores homeostasis and feeds back to repress SREBP pathway activity.

Several lines of evidence implicate SREBPs in the regulation of oncogene-driven lipogenesis. First, expression of oncogenic *PI3K* or *KRAS* is sufficient to stimulate lipogenesis in breast cancer cells through a SREBP-dependent and mTORC1-dependent mechanism (8). The vast majority of PDAC tumors contain an activating mutation in *KRAS* (9), which drives development of pancreatic intraepithelial neoplasia (PanINs) (10), and subsequent mutation of tumor suppressors results in invasive PDAC (9). Second, overexpression of the *MYC* oncogene promotes progression of *Kras*-driven PanIN to PDAC in a mouse model of pancreas cancer (11), and *MYC* cooperates with SREBPs to control lipogenesis across a number of *MYC*-dependent cancers (12). Finally, studies of the SREBP pathway in several cancers have shown a requirement for cancer cell growth, including brain, breast, colon, liver, and prostate (13–18).

Given this evidence, we hypothesized that PDAC cells activate the SREBP pathway as part of metabolic reprogramming to maintain lipid supply and support tumor growth. As such, we predicted that inhibition of the SREBP pathway should limit PDAC tumor growth. Here, we examined the requirement for the SREBP pathway and specifically its essential regulator SCAP for (1) PDAC initiation and disease progression in a genetically engineered mouse model of pancreatic cancer, (2) growth of PDAC tumor xenografts, and (3) growth of PDAC cells in culture. We demonstrate that the SREBP pathway and SCAP are required for PDAC cell survival and tumor growth, making the SREBP pathway a candidate therapeutic target for PDAC.

## MATERIALS AND METHODS

Additional information may be found in Supplementary Data, including chemical reagents, CRISPR knockout cell line generation, real-time qPCR, cell growth assays, mouse husbandry and genotyping, glucose and insulin tolerance tests, tissue membrane fractionation, protein extraction and immunoblot analysis, microarray analysis, bioinformatic analysis, and histology and photography.

### Cell Culture

Human pancreatic ductal adenocarcinoma (PDAC) cell lines Pa02c, Pa03c, Pa16c, and Pa20c were generously provided by Dr. Anirban Maitra (19,20). Pa02c and Pa03c were derived from a liver metastatic site, and Pa16c and Pa20c were derived from a primary PDAC site. Cell lines were validated by sequencing to contain *KRAS* and *TP53* mutations and verified as *Mycoplasma* negative (MycoAlert Mycoplasma Detection Kit, Lonza, LT07-701). All cells were maintained in monolayer culture at 37°C and 5% CO_2_. Wildtype cells were cultured in DMEM medium [4.5 g/L glucose, L-glutamine, and sodium pyruvate (Corning, 10-013)] supplemented with 10% (v/v) fetal bovine serum (FBS, Gibco, 10438026) and 1,000 U/mL penicillin-streptomycin (Gibco, 15140122). *SCAP* KO cells were maintained in M19 medium [(DMEM + 4.5 g/L glucose, L-glutamine, and sodium pyruvate) supplemented with 10% (v/v) FBS, 1,000 U/mL penicillin-streptomycin, 1 mM mevalonate (MilliporeSigma, M4667), 5 µg/mL cholesterol (MilliporeSigma C3045) in ethanol, and 20 µM oleic acid-albumin (MilliporeSigma O3008)]. For sterol depletion, cells were cultured as indicated in figure legends using DMEM medium (4.5 g/L glucose, L-glutamine, and sodium pyruvate) supplemented either with 10% lipoprotein-deficient serum (LPDS, Kalen 880100), 1% (v/v) FBS, or 0.5% (v/v) FBS.

### Mouse Strains

The Johns Hopkins University animal care and use program is accredited by AAALAC International, and the Johns Hopkins Institutional Animal Care and Use Committee (IACUC) reviewed and approved all mouse experimental procedures. The following mice were obtained from Jackson Laboratories: *Pdx1-Cre^+/-^* [B6.FVB-Tg(*Pdx1-Cre*)6Tuv/J] (21) (Jax #014647); *Scap^fl/fl^*(B6;129-*Scap^tm1Mbjg^*/J) (Jax #004162) (22); C57BL/6J (Jax #000664). *Kras^LSL-G12D/+^*; *Trp53^LSL-R172H/+^*(denoted as KP) mice on a congenic C57BL/6J background were generously gifted from Dr. Lei Zheng (Department of Oncology, Johns Hopkins University School of Medicine) (23). *Scap^fl/fl^* (denoted as S^fl/fl^) mice were backcrossed on to C57BL/6J background for a minimum of 6 generations. *Pdx1-Cre^+/-^*(denoted as C) mice were maintained and used as a hemizygous strain, and the *Trp53* allele was maintained as heterozygous for the entire study. All mice were maintained on a C57BL/6J background for the entire study.

### Mouse Subcutaneous Xenograft

Female athymic nude mice (Hsd:Athymic Nude-*Foxn1^nu^*, Envigo) aged 8-12 weeks old were anesthetized with isoflurane (MWI Animal Health, 501017) and injected in the subcutaneous space of each flank with 100 µL of indicated cells (5×10^5^ for Pa03c cell lines, 1×10^6^ for all other cell lines) in 50:50 (v/v) serum-free DMEM:Matrigel (Corning, 356234) for a total of two tumors per mouse. Once visible (∼7-20 days post-injection), tumors were measured along the length and width using digital calipers (Fowler) every 2-3 days. Tumor volume was calculated with the following formula: (minimum measurement^2^ x maximum measurement)/2 (24). All mice were euthanized when tumors became ulcerated or tumor burden exceeded limits set by Johns Hopkins Research Animal Resources. At euthanasia, tumors were harvested and fixed in 10% (v/v) neutral buffered formalin (Sigma, HT501128) for histologic analysis. Statistical analysis was performed in GraphPad Prism (Version 9.3.1).

### Mouse Orthotopic Xenograft

Female athymic nude mice (Hsd:Athymic Nude-*Foxn1^nu^*, Envigo) aged 8-10 weeks old were anesthetized with isoflurane and injected in the tail of the pancreas with 20 µL of indicated cells in 50:50 (v/v) serum-free DMEM:Matrigel. Briefly, mice were placed in right lateral recumbency, and a 5-7 mm transverse incision was made on the left flank skin and underlying body wall caudal to the ribs. The tail of the pancreas was exteriorized, and 20 µL of cell suspension containing 50,000 cells was injected using a Hamilton syringe with a 30-gauge needle. Successful injection was confirmed by the presence of swelling within the pancreatic tissue. The body wall and skin were sutured closed in a simple continuous and simple interrupted pattern, respectively, using 4-0 absorbable sutures, and the skin edges sealed with tissue glue (3M VetBone). All mice received a 1 mg/kg subcutaneous injection of Burprenorphine-SR Lab (1 mg/mL, ZooPharm) prior to recovery with an additional dose given as needed every 48 hours. Mice were weighed at surgery and every 5-7 days post-surgery. On day 30 post-surgery, all mice were euthanized. Tumors were harvested and fixed in 10% (v/v) neutral buffered formalin for histologic analysis. Statistical analysis was performed using GraphPad Prism (Version 9.3.1).

### Survival Study Histologic Analysis

Mice were bred to obtain the following cohorts: <u>KPC</u> (*Kras^LSL-G12D/+^*; *Trp53^LSL-R172H/+^*; *Pdx1-Cre^+/-^*); KPCS^fl/+^ (*Kras^LSL-G12D/+^*; *Trp53^LSL-R172H/+^*; *Pdx1-Cre^+/-^*; *Scap^fl/+^*); KPCS^fl/fl^ (*Kras^LSL-G12D/+^*; *Trp53^LSL-R172H/+^*; *Pdx1-Cre^+/-^*; *Scap^fl/fl^*); <u>C</u> (*Pdx1-Cre^+/-^)*; CS^fl/+^ (*Pdx1-Cre^+/-^*; *Scap^fl/+^*); CS^fl/fl^ (*Pdx1-Cre^+/-^*; *Scap^fl/fl^*). All mice with appropriate genotypes and of both sexes were initially included in the study. Mice were allowed to age and were euthanized upon any of the following clinical signs of morbidity: labored breathing, abdominal distension, hunched/lethargic, poor body condition, large (exceeded 2 cm in any direction) and/or ulcerated external masses that affected clinical status, moribund status, or other clinical disease where euthanasia was recommended by JHU Research Animal Resources. Most control cohorts (C, CS^fl/+,^ and CS^fl/fl^) were euthanized when they reached at least 450 days of age to terminate the study. Full necropsy/gross examination was performed, and more than 30 tissues fixed in 10% neutral buffered formalin (25). Exclusion criteria included the following: death at <60 days of age, severe post-mortem autolysis/decomposition or desiccation at gross examination, or severe post-mortem autolysis at histologic examination that precluded any reasonable interpretation of majority of the tissues. Due to the nature of the genetic mutations and resulting phenotypes, investigators could not be completely blinded to the mouse genotype during analysis. Histopathologic analysis of all tissues was performed by a boarded veterinary pathologist to confirm cause of death or euthanasia as well as assess the presence of pancreatic intraepithelial neoplasia (PanINs) (10) and invasive PDAC. PanIN grading was not applied in this study. Analysis of the pancreas specifically was confirmed by an additional boarded pathologist. For survival analysis, the adverse event was strictly defined as death or euthanasia secondary to invasive PDAC. Death secondary to PDAC was assigned if euthanasia or death was due to any combination of the following: presence of obstructive PDAC, presence of PDAC metastases to any other organ, presence of malignant abdominal effusion, and/or carcinomatosis. All other causes of death/euthanasia (**Table 1** and **Table S1**) were classified based on clinical, gross and histopathological findings.

**TABLE 1.** Causes of death in KPC mouse cohorts.

| Cause of Death/Euthanasia | KPC | KPCS<sup>fl/+</sup> | KPCS<sup>fl/fl</sup> |
| --- | --- | --- | --- |
| PDAC | 8/18 (44.4%) | 4/32 (12.5%) | 0/16 (0%) |
| Hematopoietic neoplasia (e.g. lymphoma) | 1/18 (5.6%) | 7/32 (21.9%) | 1/16 (6.3%) |
| Large/ulcerated epidermal tumors (facial or perineal) | 3/18 (16.7%) | 5/32 (15.6%) | 3/16 (18.8%) |
| Thyroid or salivary gland tumor | 2/18 (11.1%) | 6/32 (18.8%) | 1/16 (6.3%) |
| Other/miscellaneous | 2/18 (11.1%) | 4/32 (12.5%) | 8/16 (50%) |
| Unknown | 2/18 (11.1%) | 6/32 (18.8%) | 3/16 (18.8%) |
| **Total** | **18/18 (100%)** | **32/32 (100%)** | **16/16 (100%)** |

### Survival Study Statistical Analysis

Cumulative incidence of death due to PDAC was estimated using the competing risk method, where death due to other causes was considered as a competing event, and Gray’s test was performed for comparisons between genotype groups. The presence of PDAC was examined at the time of death of a mouse, but the exact time of developing PDAC was not observed. Therefore, an interval-censored survival analysis was employed to characterize time to development of PDAC. Specifically, survival probabilities were estimated based on the non-parametric maximum likelihood method implemented in the “icfit” function in the R package “interval” (Version 1.1-0.8) (26). Permutation test was performed to compare the time to PDAC development between genotype groups. All tests were two-sided and p value <0.05 was considered to indicate statistical significance. The analysis was carried out using R software (version 4.2.1).

## Data Availability

The data generated in this study are publicly available in Gene Expression Omnibus (GEO) at GSE235100.

## RESULTS

### Scap is not required for development or function of the mouse pancreas

PDAC tumors are characterized by a dense, fibrotic stroma that impedes vascularization, resulting in limited oxygen and nutrient delivery to tumor cells (27). Given that an abundant supply of lipids is required for PDAC tumor cells to proliferate and survive, we hypothesized that SREBP pathway activity is required for PDAC tumor growth. SCAP is required for the activation of both SREBP-1 and SREBP-2, and loss of *SCAP* abolishes SREBP pathway activity (7). To test whether the SREBP pathway is required for the development and progression of PDAC, we set out to test whether loss of *Scap* impacts disease in the well-established KPC genetically engineered mouse model (GEMM) of PDAC, *LSL-<u>K</u>ras^G12D/+^, LSL-Tr<u>p</u>53^R172H/+^, Pdx1-<u>C</u>re* (KPC) mice (28).

KPC mice offer the advantage of an accelerated tumor model that pathologically recapitulates many aspects of human disease (28). This GEMM maintains an intact immune system as well as spontaneous tumor development secondary to specific gene mutations. KPC mice contain mutant alleles of both *Kras* and *Trp53,* whose expression is activated by Cre-mediated removal of lox-STOP-lox cassettes (LSL). Cre expression is driven by the pancreas-specific *Pdx-1* promoter. Importantly, the requirement of *Scap* for mouse pancreas development and function had not been examined. So, we first performed experiments to determine the role of *Scap* in mouse pancreatic function using mice carrying *Pdx1-<u>C</u>re* and an existing loss-of-function, floxed *Scap* allele (22).

Compared to wildtype, *Pdx1-Cre* only (C), and floxed *Scap* (S^fl/fl^) mice, CS^fl/fl^ mice showed reduced levels of Scap protein in the pancreas, but no differences in body weight (**Supplementary Fig. 1A-B**). Male and female CS^fl/fl^ mice showed no evidence of pancreatic endocrine dysfunction as assessed by glucose and insulin tolerance tests (**Supplementary Fig. 1C-D**) and had normal pancreas architecture histologically (**Supplementary Fig. 1E**). Thus, we concluded that *Scap* is not required for development and function of the mouse pancreas, and pancreas-specific deletion of *Scap* did not cause gross metabolic abnormalities. Based on these results, we do not expect defects in pancreas function to confound interpretation of KPC mouse model experiments.

### SCAP supports PDAC development and loss of SCAP prolongs survival in the KPC mouse model of pancreas cancer

To test the requirement of *Scap* for the development of PDAC in the KPC model, we generated KPC mice with both heterozygous (KPCS^fl/+^) and homozygous (KPCS^fl/fl^) loss of *Scap* as well as appropriate control mice (**Supplementary Fig. 2A-B**). Mouse cohorts were aged until death, and cause of death was determined **(Table 1, Supplementary Fig. 2C**). To determine the requirement of *Scap* for PDAC in KPC mice, we defined the adverse event as death due to PDAC disease and considered death due to other causes as a competing event. When compared to KPC mice, KPCS^fl/+^ mice had a significantly prolonged survival (**Fig. 1A**). Likewise, KPCS^fl/+^ mice also showed significantly prolonged survival compared to KPC mice when we treated death due to other causes as a censoring event (**Supplementary Fig. 2D**). KPCS^fl/fl^ mice did not live as long as KPCS^fl/+^ or KPC mice, which prevented evaluation of homozygous loss of *Scap* on PDAC survival (**Fig. 1A, Supplementary Fig. 2C-D**). This observation suggests that an unidentified metabolic phenotype likely exists in KPCS^fl/fl^ mice. None of the control cohorts (C, CS^fl/+^, and CS^fl/fl^) reached the adverse event (**Supplementary Fig. 2C**). Overall, death due to PDAC was reduced in mice with a heterozygous loss of *Scap*, and completely absent in mice with a homozygous loss of *Scap* (**Fig. 1B**).

**FIGURE 1.**
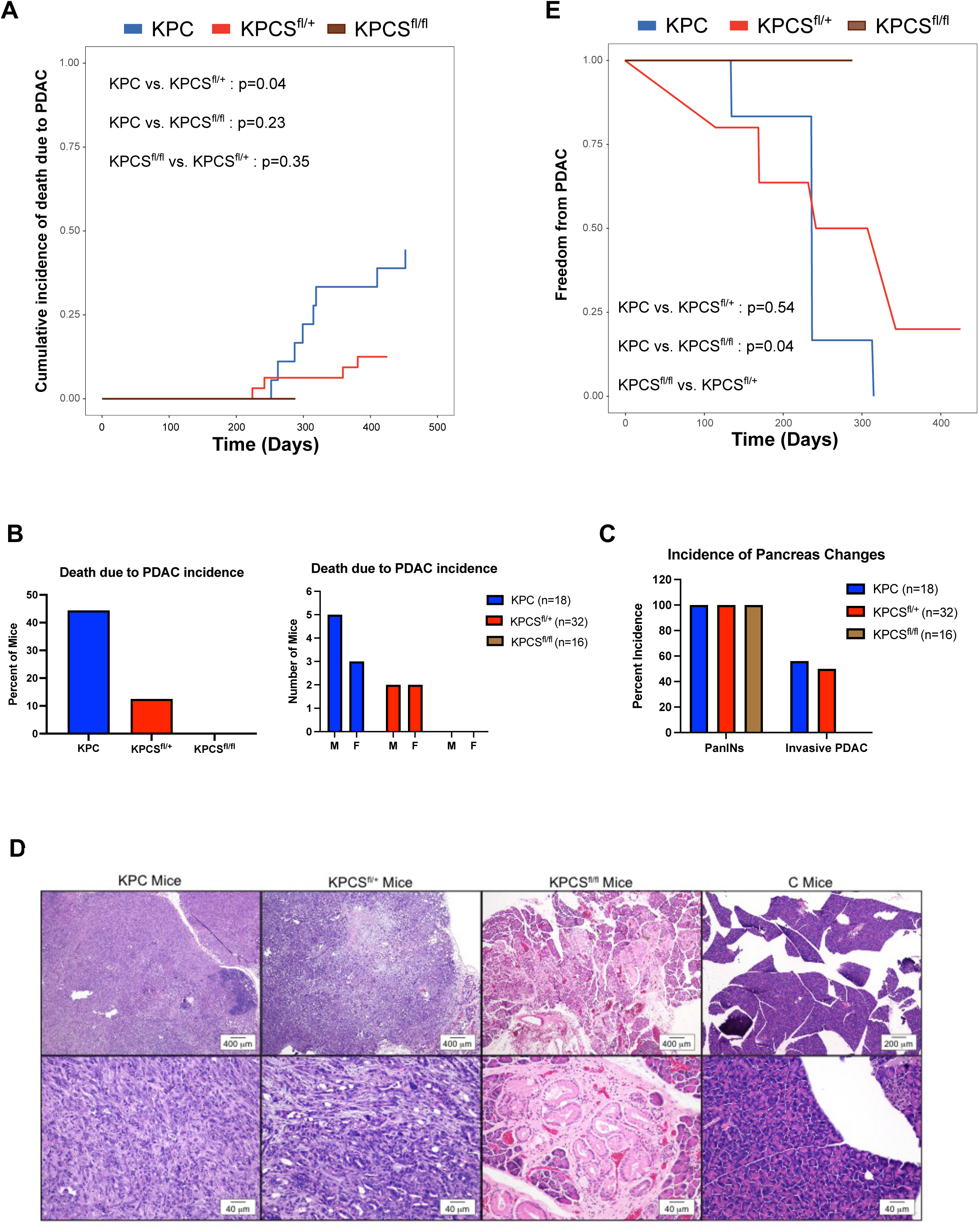
*SCAP* supports PDAC development and loss of *SCAP* prolongs survival in the KPC mouse model of pancreas cancer. **A)** Mice of the indicated genotype were aged until death. Figure shows the cumulative incidence curve of death due to PDAC for each genotype group. For the analysis, we considered death due to other causes (**i.e.**, not due to PDAC) as a competing event. Gray’s test was used for pairwise comparisons. **B)** Incidence of adverse event (death due to PDAC) in KPC cohorts from A either pooled (left figure) or stratified by sex (right figure). **C)** Incidence of PanINs and invasive PDAC identified histologically in KPC cohorts from A. **D)** Representative H&E-stained sections of formalin fixed pancreas tissue from KPC, KPCS^fl/+^ and KPCS^fl/fl^ cohorts in A showing a magnification (2-4X) image (upper row) and a higher magnification (20X) image (bottom row) of the changes in the pancreas. The C mouse is used as a control. **E)** Figure shows the survival curves by genotype group. Mice of the indicated genotypes were aged until death at which time histology was performed to determine the presence of PanINs and invasive PDAC. For this analysis, we treated time to invasive PDAC as interval censored data. The permutation test was used for pairwise comparisons.

As the development of PDAC has known progressive histologic features (29), we histologically evaluated the presence of precursor PanINs and invasive ductal adenocarcinoma in mouse cohorts. All mice in the KPC cohorts developed PanINs (**Fig. 1C-D, Table S1**), whereas none of the control mice lacking *Kras* and *Trp53* activation developed these precursor lesions. Additionally, half of KPC and KPCS^fl/+^ mice also developed invasive ductal adenocarcinoma (**Fig. 1C, Table S1**). Strikingly, none of the KPCS^fl/fl^ mice developed invasive ductal adenocarcinoma (**Fig. 1C-D, Table S1**). To test whether loss of *Scap* affected progression to PDAC as indicated by the presence of invasive carcinoma, we treated the time to PDAC as an interval censored event.

Statistical analysis showed that KPCS^fl/fl^ mice had a significant delay in the onset of invasive carcinoma compared to both KPC and KPCS^fl/+^ cohorts (**Fig. 1E**). There was no difference in time to development of invasive carcinoma between KPC and KPCS^fl/+^ cohorts (**Fig. 1E**). Given that activated *Kras* drives PanIN formation (21), these results suggest that loss of *Scap* has minimal impact on early *Kras*-dependent carcinogenesis in mouse PDAC. Taken together, these *in vivo* experiments demonstrate that *Scap* is required for the progression to PDAC and suggest that the prolonged survival of KPCS^fl/+^ mice is due in part to a delay in full disease onset. Having demonstrated a key role for *Scap* in the pathophysiology of PDAC in KPC mice, we shifted our focus to understanding the mechanistic requirement for the SREBP pathway using human PDAC cell lines.

### SCAP is required for human PDAC tumor growth in mouse xenograft models

To investigate the requirement of the SREBP pathway for human PDAC tumor growth, we examined multiple human cell lines from both primary PDAC tumors (Pa16c and Pa20c) and liver metastases (Pa02c and Pa03c). We created stable *SCAP* knockout (KO) cell lines for each using CRISPR technology. *SCAP* KO cells were confirmed by sequencing and had no detectable SCAP protein (**Supplementary Fig. 3**). To extend our results from the KPC mice (**Fig. 1**), we tested whether *SCAP* is required for human PDAC tumor establishment and progression in mouse xenograft models. Compared to wildtype human Pa03c cells, *SCAP* KO Pa03c cells showed significantly decreased tumor growth in subcutaneous xenografts in nude mice (**Fig. 2A-B**). Similarly, growth of Pa16c and Pa20c subcutaneous xenografts required *SCAP* (**Fig. 2D-E, Supplementary Fig. 4A-B**), while growth of Pa02c xenografts was inhibited but not completely as seen with the other three cell lines (**Supplementary Fig. 4C-D**). On histologic evaluation, wildtype tumors were highly cellular with a trabecular or ductal pattern, while *SCAP* KO tumors had significantly decreased cellularity, loss of trabecular organization, increased cellular vacuolation (indicative of cellular degeneration), and increased interweaving fibrous tissue (**Fig. 2C, F, Supplementary Fig. 4E**).

**FIGURE 2.**
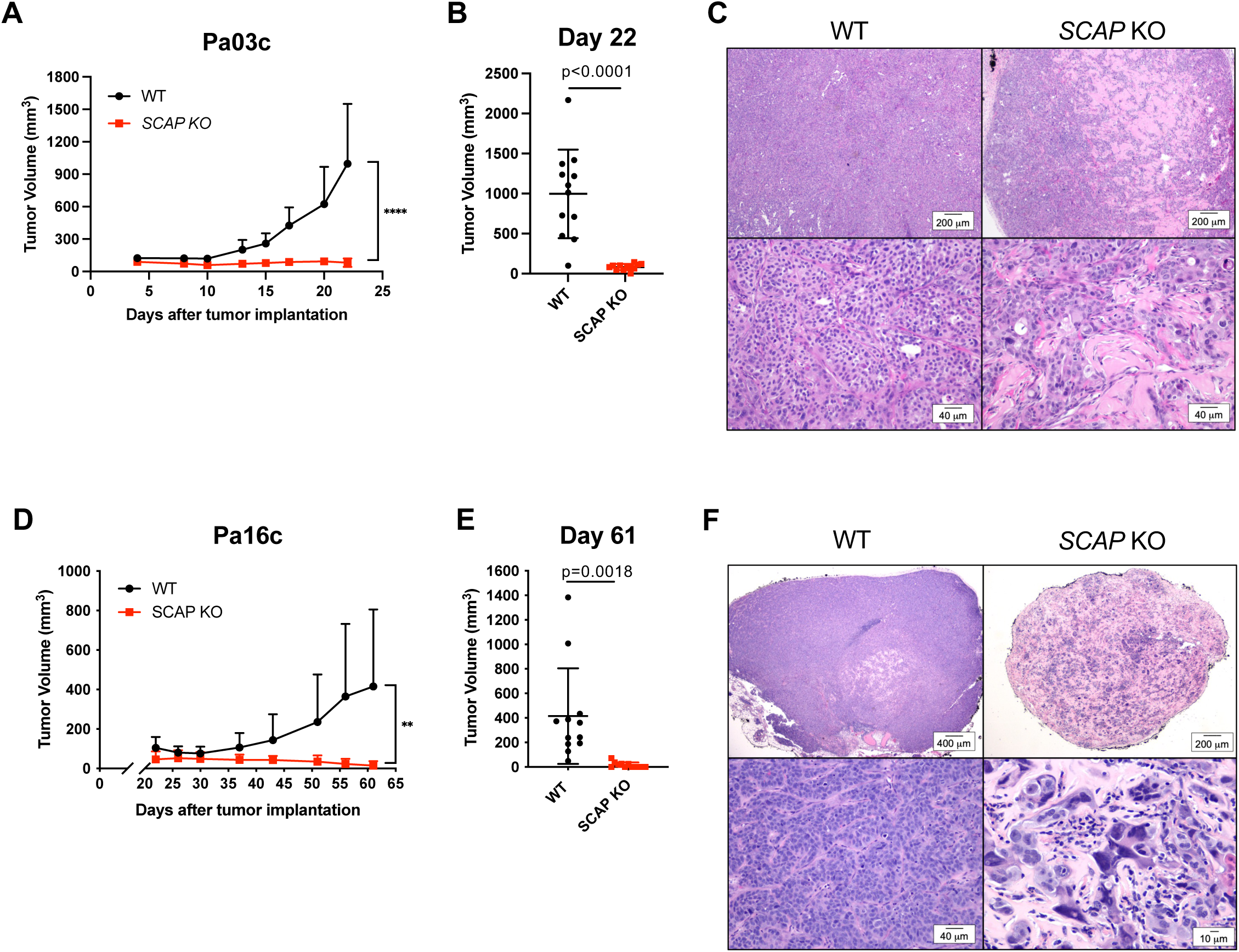
– *SCAP* is required for human PDAC tumor growth in mouse subcutaneous xenograft models. **A)** Nude mice were subcutaneously injected with 5×10^5^ Pa03c cells in both flanks (two tumors per mouse). Once visible, tumors were measured, and volume calculated. Each group contained 6 mice. Error bar denotes standard deviation. Statistical significance was determined using student’s t-test. P values are indicated: < 0.05 (*); < 0.01 (**); < 0.001 (***); < 0.0001 (****), not significant (ns). **B)** Individual tumor volumes at day 22, n=12 tumors per group. Statistical significance was determined using student’s t-test. **C)** Representative hematoxylin and eosin (H&E) stained sections of formalin fixed tumor tissue from the mice in A showing a low magnification (4X) image (upper row) and a higher magnification (20X) image (bottom row) of tumor sections. **D)** Nude mice were subcutaneously injected with 5×10^5^ Pa16c cells in both flanks (two tumors per mouse) as in A-C. Each group contained 6 mice. Error bar denotes standard deviation. Statistical significance was determined using student’s t-test. P values are indicated: < 0.05 (*); < 0.01 (**); < 0.001 (***); < 0.0001 (****), not significant (ns). **E)** Individual tumor volumes at day 61, n=12 tumors per group. Statistical significance was determined using student’s t-test. **F)** Representative hematoxylin and eosin (H&E) stained sections of formalin fixed tumor tissue from the mice in D showing a low magnification (2-4X) image (upper row) and a higher magnification (20-40X) image (bottom row) of tumor sections.

To test whether growth of human PDAC xenografts requires *SCAP* in a model that more accurately represents the tumor microenvironment, we utilized an orthotopic xenograft model. Wildtype human Pa03c and *SCAP* KO cells were injected into the tail of the pancreas in nude mice, and mice were examined for tumors after 30 days. Pa03c *SCAP* KO cell tumors were significantly smaller than wildtype Pa03c cell tumors (**Fig. 3A-B**).

**FIGURE 3.**
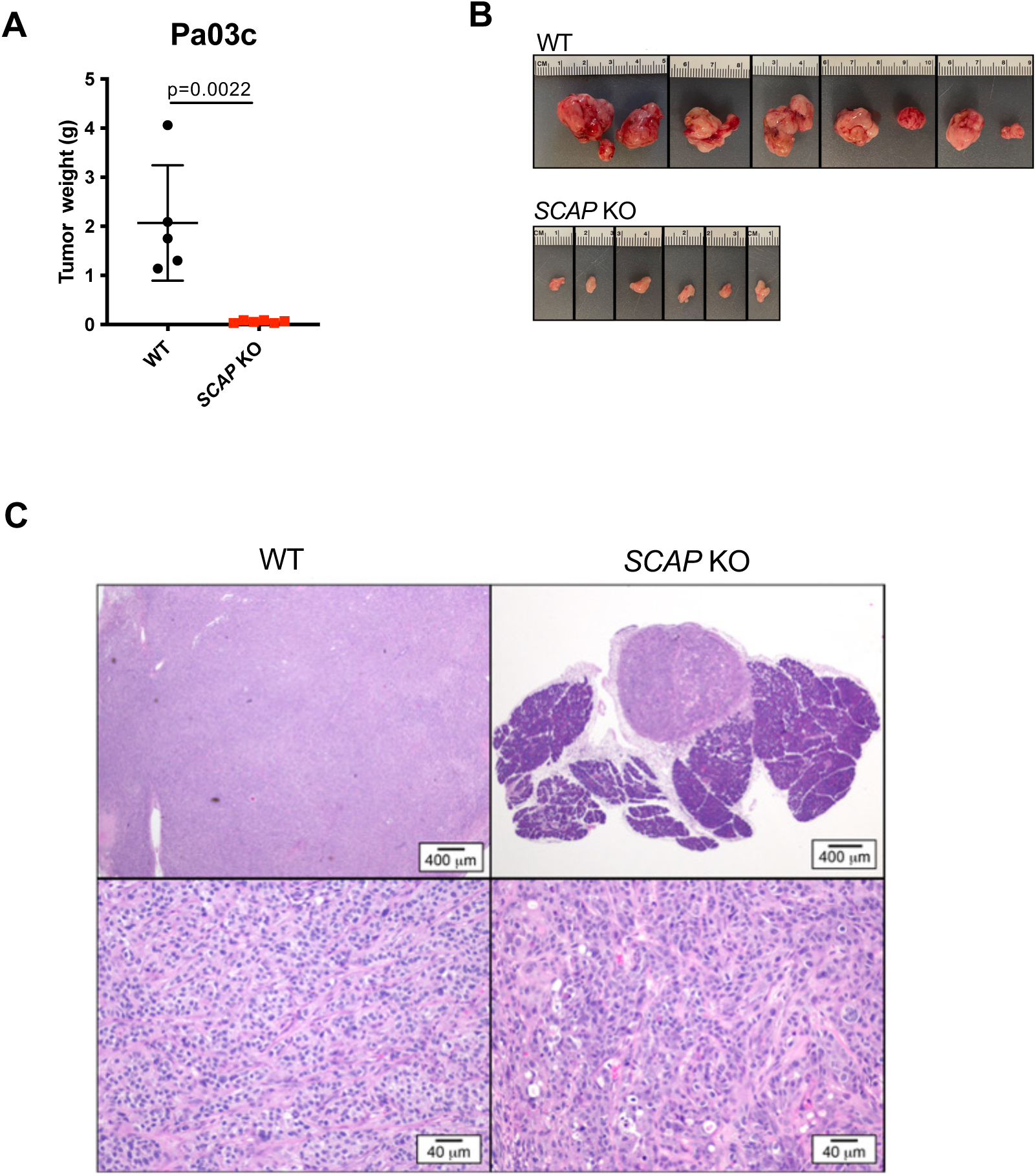
*SCAP* is required for human PDAC tumor growth in mouse orthotopic xenograft models. **A)** Nude mice were orthotopically injected with 5×10^4^ Pa03c cells in the tail of the pancreas (one tumor per mouse). Mice were euthanized on day 30 post-surgery and tumors weighed. The wildtype group contained 5 mice, and the *SCAP* KO group contained 6 mice. Error bar denotes standard deviation. Statistical significance was determined using student’s t-test. **B)** Photograph images of all harvested tumors from A. **C)** Representative H&E-stained sections of formalin fixed tumor tissue from the mice in A showing a low magnification (2X) image (upper row) and a higher magnification (20X) image (bottom row) of the tumors.

Microscopically, wildtype Pa03c cell tumors from the pancreas orthotopic xenograft model showed an organized trabecular pattern similar to the subcutaneous xenograft tumors, while *SCAP* KO cell tumors showed a distinct loss of architecture with more cells undergoing single cell necrosis (**Fig. 3C**). Together, these xenograft models demonstrate that *SCAP* is required for human PDAC tumor growth both subcutaneously and in the pancreas. Importantly, these data parallel those from the KPC mouse study and confirm that SREBP pathway activity is an absolute requirement for PDAC tumor growth.

*SCAP is required for PDAC cell growth under low serum and low lipid conditions* PDAC tumors are poorly vascularized, which limits nutrient delivery to tumor cells (27). To investigate how nutrient-poor conditions impact global gene expression in human PDAC cells, we assayed gene expression comparing two serum conditions and performed gene set enrichment analysis. Human Pa03c cells were cultured in medium containing 0.5% FBS and compared to cells cultured in medium containing 10% FBS. We identified 187 genes whose expression was increased and 158 genes whose expression was decreased (**Table S2**). Differentially expressed genes were analyzed against four separate cellular pathway databases: Reactome, Hallmark, WikiPathways, and KEGG. Notably, SREBP target genes and related biochemical pathways were the most strongly enriched among the genes upregulated in 0.5% FBS as indicated by the fact that expression analysis identified the SREBP pathway, cholesterol metabolism, and terpenoid/steroid synthesis in each of the databases (**Fig. 4A**). Genes with decreased expression in 0.5% FBS compared to 10% FBS were broadly associated with cell metabolism and growth **(Table S2).** Thus, SREBP target gene expression is broadly upregulated in PDAC cells under low serum conditions, demonstrating that PDAC cells activate the SREBP pathway in a nutrient-poor environment.

**FIGURE 4.**
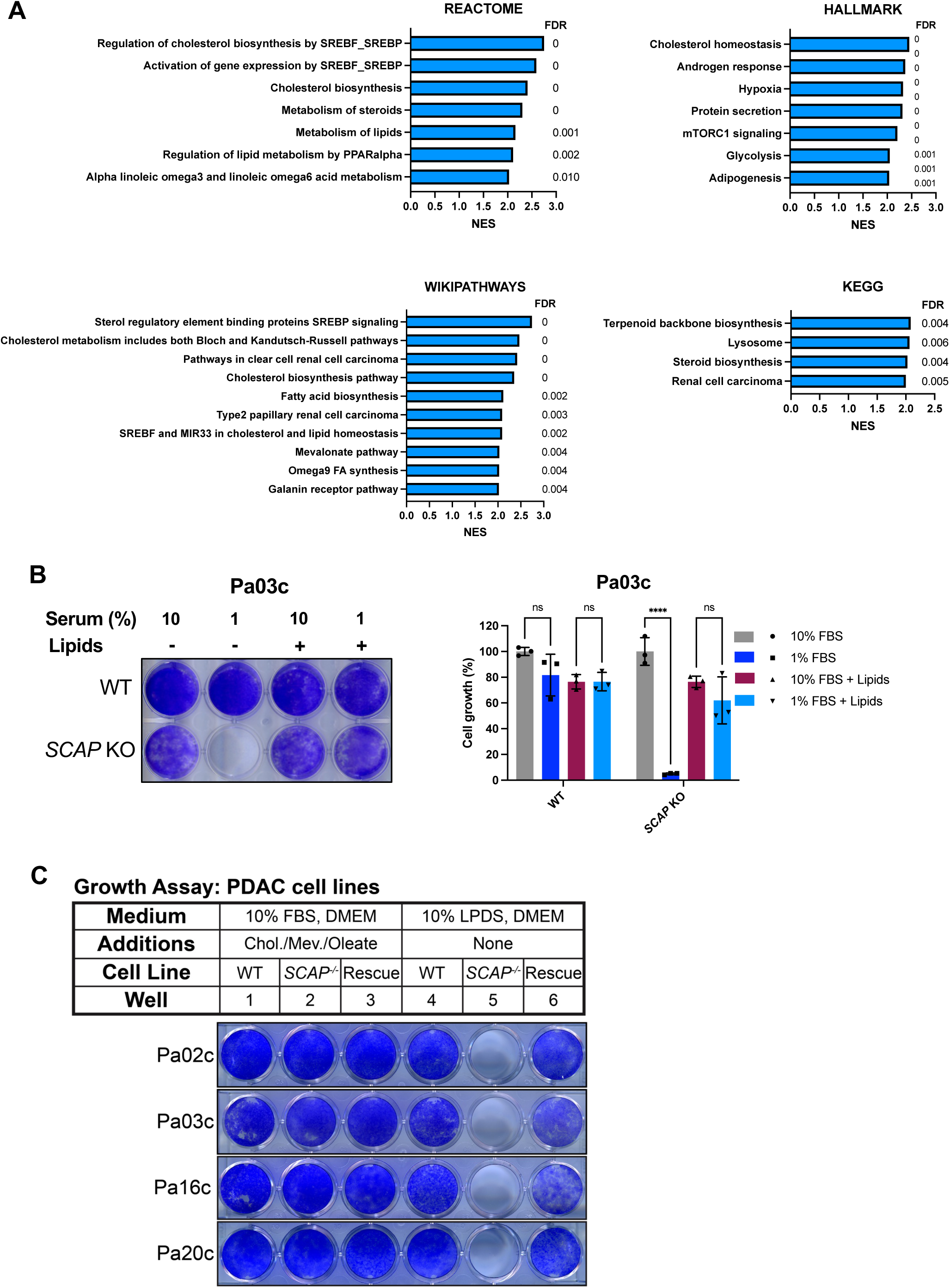
*SCAP* is required for PDAC cell growth under low serum and low lipid conditions. **A)** Pa03c cells were grown for 16 hours in medium containing 10% FBS or 0.5% FBS. mRNA expression was determined using microarray, and differentially expressed genes were identified **(Table S2)**. Bioinformatic gene set enrichment analysis was performed against 4 common pathway datasets. Results are shown using a normalized enrichment score (NES) > 2.0. FDR denotes false discovery rate. **B)** Wildtype and *SCAP* KO Pa03c cells were grown for 7 days in medium containing 10% FBS or 1% FBS in the absence or presence of lipid supplements (1 mM mevalonate, 5 µg/mL cholesterol in ethanol, 20 µM sodium oleate, 50 µg/mL LDL). Media were changed every 3 days. Plates were stained with crystal violet as a measure of cell proliferation. Quantification of crystal violet staining is shown (n = 3 per group). For each cell line, growth was normalized to the 10% FBS condition. Statistical significance was determined using two-way ANOVA and Tukey’s HSD test. P values are indicated: < 0.05 (*); < 0.01 (**); < 0.001 (***); < 0.0001 (****), not significant (ns). Error bar denotes standard deviation. **C)** Cell growth assay of PDAC cell lines. WT, *SCAP* KO, and *SCAP* KO rescued cell lines were cultured in either 10% FBS supplemented with cholesterol (5 µg/mL), mevalonate (1 mM) and oleate-albumin (20 µM) or 10% LPDS with no additions for 7 days. Plates were stained with crystal violet.

To investigate what underlies the requirement for *SCAP* in the PDAC mouse tumor models (Fig. 1-3), we examined the requirement of *SCAP* for human PDAC cell growth *in vitro*. Given our gene expression results, we hypothesized that *SCAP* functions to upregulate SREBP in a homeostatic response to low serum conditions. We assayed growth of wildtype Pa03c and *SCAP* KO cells in medium containing either 10% FBS or 1% FBS to compare nutrient-rich and nutrient-poor environments. Wildtype and *SCAP* KO Pa03c cells grew similarly in medium containing 10% FBS (**Fig. 4B**). Remarkably, *SCAP* KO Pa03c cells failed to grow in 1% FBS, indicating an essential requirement for SREBP activation under nutrient-poor conditions (**Fig. 4B**). The growth defect in *SCAP* KO Pa03c cells was rescued by the addition of lipids (cholesterol, mevalonate, oleic acid, and LDL), indicating that *SCAP* KO cells failed to grow due to lack of SREBP-dependent lipid supply (**Fig. 4B**). Similar results were observed with Pa02c and Pa20c cell lines (**Supplementary Fig. 5**). FBS contains multiple growth factors and nutrients other than lipids that may contribute to the requirement of *SCAP* for growth in 1% serum. To test whether *SCAP* is specifically required for cell growth under low-lipid conditions, we cultured cells in either 10% lipoprotein-deficient serum (LPDS) or lipid-rich medium that contains 10% FBS and supplemental cholesterol, mevalonate and oleate. All four PDAC cell lines lacking *SCAP* failed to grow in LPDS-containing medium compared to the wildtype parental cell line, but *SCAP* KO cells grew as well as wildtype in lipid-rich medium (**Fig. 4C**). To confirm that the growth defect in *SCAP* KO cells was due to targeted mutation of *SCAP*, we generated stable *SCAP* KO cell lines expressing *SCAP.* Reintroduction of *SCAP* restored cell growth in LPDS-containing medium (**Fig. 4C**), indicating that the lipid-dependent growth defects were due to loss of *SCAP*. Collectively, these data demonstrate that *SCAP* is conditionally essential for PDAC cell growth, required under lipid-poor but not lipid-rich conditions.

### SREBP pathway is required for PDAC cell growth in low serum conditions

SCAP functions to activate SREBPs in order to maintain cellular lipid homeostasis (7). Given the requirement of *SCAP* for low serum growth, we next examined regulation of SREBPs under these conditions. Thus far, we have used genetic inhibition of *SCAP* to inactivate SREBPs, so we employed two different chemical inhibitors as an independent test of the requirement for the SREBP pathway. Human Pa03c cells were grown in medium containing either 10% or 1% FBS, and the requirement for SREBP pathway activity was tested using the Site-1 protease inhibitor, PF-429242, which prevents proteolytic activation of SREBPs (30). Pa03c and Pa16c cells showed a sharp dose-dependent decrease in cell growth when treated with PF-429242 in 1% FBS (**Fig. 5A**). Growth inhibition by PF-429242 was low serum-dependent as PF-429242 only affected cell growth in 10% FBS at concentrations ≥ 10 µM. Given our gene expression data (**Fig. 4A**), we next examined whether SREBPs were proteolytically activated in PDAC cells under low serum conditions. The nuclear form of both SREBP-1 and SREBP-2 increased in PDAC cells after growth in medium containing 1% FBS (**Fig. 5B**). SREBP-1 and SREBP-2 cleavage were markedly inhibited upon treatment with the Site-1 protease inhibitor PF-429242 in 1% FBS, confirming that PF-429242 inhibits SREBP activation (**Fig. 5B**). Consistent with these results, mRNA levels of SREBP-1 target genes (*SCD*; *fatty acid synthase*, *FASN*; *insulin-induced gene 1*; *INSIG1*) and SREBP-2 target genes (*HMGCR*; *HMG-CoA synthase 1, HMGCS1*; *LDLR*) and their corresponding protein products were upregulated in medium containing 1% FBS (**Fig. 5C-D**). As expected, SREBP target gene mRNAs and proteins sharply decreased after treatment with PF-429242 in 1% FBS (**Fig. 5C-D**). Similar results were observed with Pa02c and Pa20c cell lines (**Supplementary Fig. 6A-C**). These results demonstrate that the SREBP pathway is activated in PDAC cells under low serum conditions and that pathway activity is conditionally essential for growth in low serum conditions *in vitro*.

**FIGURE 5.**
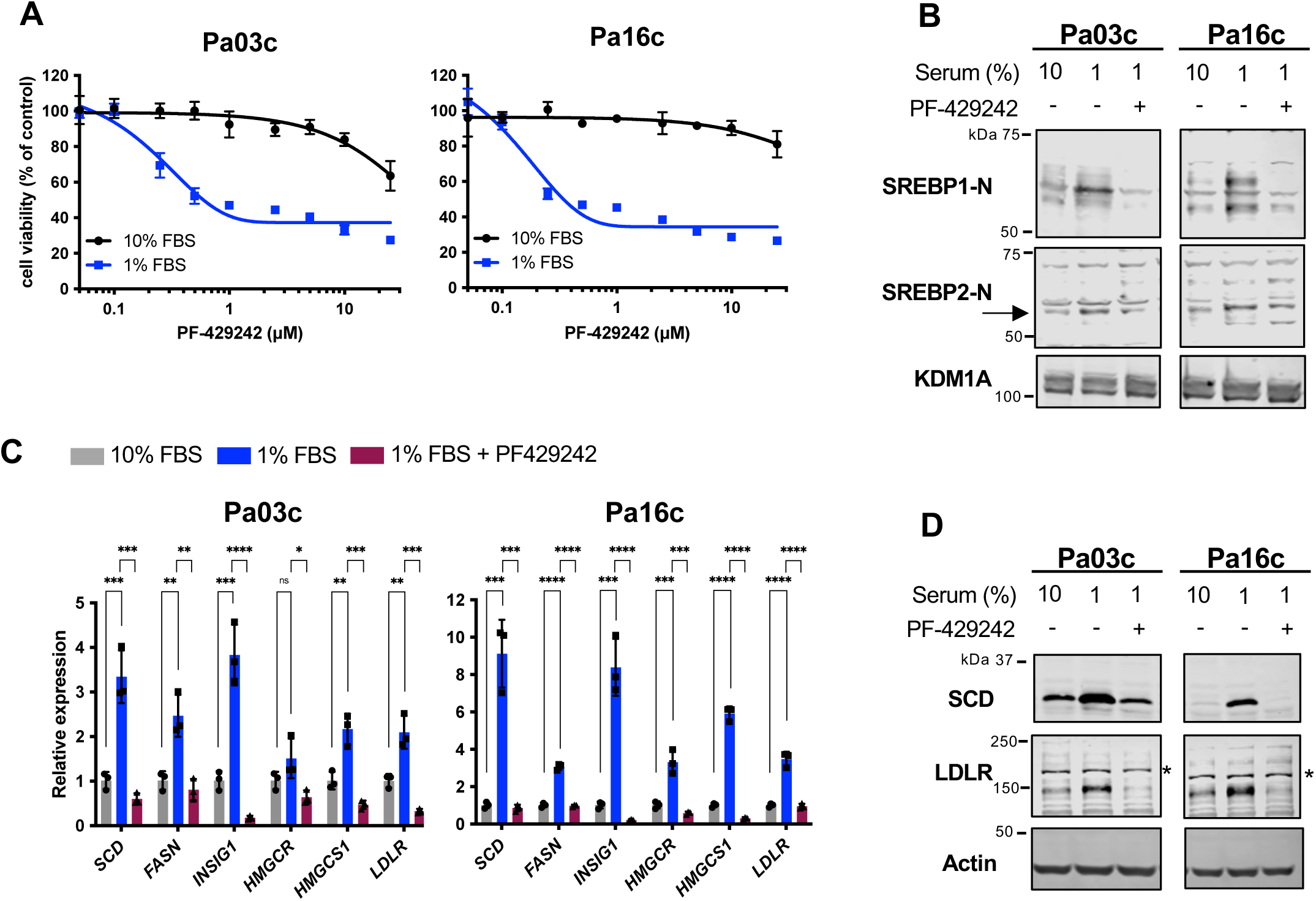
Site-1 protease activation of the SREBP pathway is required for PDAC cell growth in low serum conditions. **A)** PDAC cells (Pa03c and Pa16c) were cultured in either 10% or 1% FBS with indicated concentrations of the Site-1 protease inhibitor PF-429242 for 72 hours, and cell growth was determined using a MTS assay with data normalized to 10% FBS untreated cells. Data are representative of 2 biological replicates with 3 technical replicates for each biological replicate. Error bar denotes standard deviation. **B)** Immunoblot analysis of nuclear extracts from human Pa03c and Pa16c cells cultured for 16 hours in either 10% FBS, 1% FBS, or 1% FBS containing PF-429242 (10 µM). Blots were probed for either SREBP1-N or SREBP2-N (arrow), and lysine-specific histone demethylase 1A (KDM1A) served as a loading control. Result is representative of 2 biological replicates. **C)** Pa03c and Pa16c cells were cultured under the same conditions as in B), and quantitative real-time PCR was performed for target genes of SREBP1 (*SCD*, *FASN*, *INSIG1*) and SREBP2 (*HMGCR*, *HMGCS1*, *LDLR*). *GAPDH* served as a control. Data are representative of 2 biological replicates with 3 technical replicates for each biological replicate. Error bar denotes standard deviation. Statistical significance was determined using one-way ANOVA and Tukey’s HSD test. P values are indicated: < 0.05 (*); < 0.01 (**); < 0.001 (***); < 0.0001 (****), not significant (ns). **D)** Immunoblot analysis of whole cell lysates from Pa03c and Pa16c cells cultured as in B) for indicated SREBP target protein expression. The asterisk indicates a non-specific band present in Pa03c and Pa16c cell lines. Actin served as a loading control. Result is representative of 2 biological replicates.

The oxysterol 25-hydroxycholesterol (25-HC) inhibits SREBP transport to the Golgi and prevents its proteolytic activation (31). We next tested the effects of 25-HC on PDAC cell growth and SREBP pathway activation in low serum conditions. Pa03c and Pa16c cells showed dose-dependent growth inhibition by 25-HC in 1% FBS containing medium (**Fig. 6A**). Growth inhibition was blunted in medium containing 10% FBS, indicating that the effects of 25-HC were serum-dependent. Compared to PF-429242 (**Fig. 5A**), 25-HC showed less serum dependence in Pa03c cells, possibly due to additional effects of 25-HC which is also a ligand for the nuclear receptor LXR (32). The active nuclear forms of SREBP-1 and SREBP-2 were upregulated in 1% FBS growth medium, and activation was inhibited upon treatment with 25-HC in 1% FBS, as expected (**Fig. 6B**). The effects on SREBP target gene expression paralleled those on the SREBP nuclear forms. Expression of target gene mRNA and protein was upregulated in medium containing 1% FBS compared to 10% FBS, and addition of 25-HC blocked upregulation (**Fig. 6C-D**). Similar results were observed with Pa02c and Pa20c cell lines (**Supplementary Fig. 7A-C**). Together, these data demonstrate that expression of SREBP target genes and their protein products are upregulated under low serum conditions due to SREBP activation through the classical pathway that requires ER-to-Golgi transport and the Site-1 protease. Notably, pharmacological inhibition of the SREBP pathway decreased PDAC cell growth under low serum conditions that we hypothesize mimics the tumor microenvironment.

**FIGURE 6.**
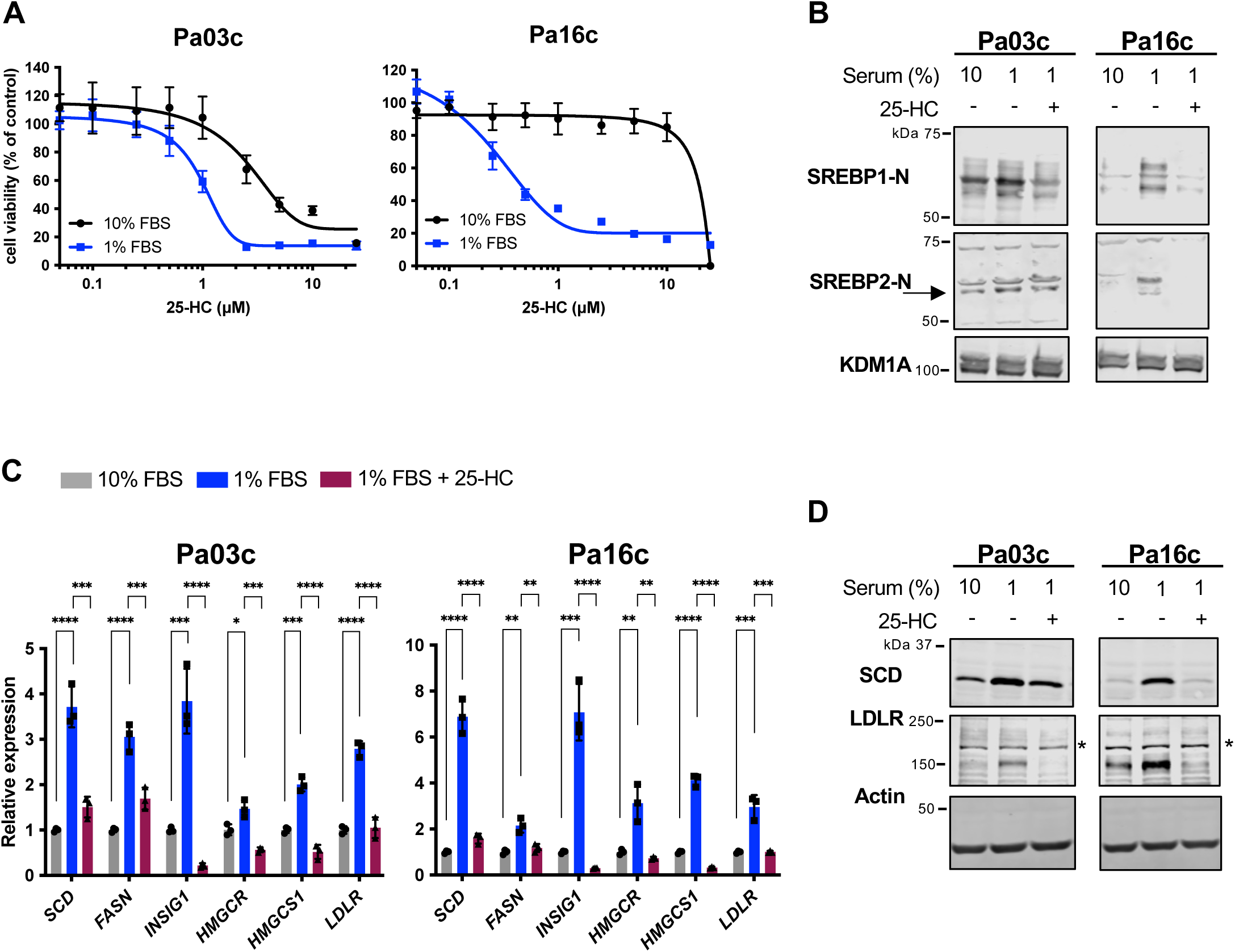
SREBP pathway activation is required for PDAC cell growth in low serum conditions. **A)** PDAC cells (Pa03c and Pa16c) were cultured in either 10% or 1% FBS with indicated concentrations of 25-hydroxycholesterol (25-HC) for 72 hours, and cell growth was determined using a MTS assay with data normalized to 10% FBS untreated cells. Data are representative of 2 biological replicates with 3 technical replicates for each biological replicate. Error bar denotes standard deviation. **B)** Immunoblot analysis of nuclear extracts from human Pa03c and Pa16c cells cultured in either 10% FBS, 1% FBS, or 1% FBS containing 25-hydroxycholesterol (2.5 µM) for 16 hours. Blots were probed for either SREBP1-N or SREBP2-N (arrow), and lysine-specific histone demethylase 1A (KDM1A) served as a loading control. Result is representative of 2 biological replicates. **C)** Pa03c and Pa16c cells were cultured under the same conditions as in B), and quantitative real-time PCR was performed for target genes of SREBP1 (*SCD*, *FASN*, *INSIG1*) and SREBP2 (*HMGCR*, *HMGCS1*, *LDLR*). *GAPDH* served as a control. Data are representative of 2 biological replicates with 3 technical replicates for each biological replicate. Error bar denotes standard deviation. Statistical significance was determined using one-way ANOVA and Tukey’s HSD test. P values are indicated: < 0.05 (*); < 0.01 (**); < 0.001 (***); < 0.0001 (****), not significant (ns). **D)** Immunoblot analysis of whole cell lysates from Pa03c and Pa16c cells cultured as in B) for indicated SREBP target protein expression. The asterisk indicates a non-specific band present in Pa03c and Pa16c cell lines. Actin served as a loading control. Result is representative of 2 biological replicates.

### Inhibition of both SREBP-1 and SREBP-2 is required to prevent PDAC cell and tumor growth

SCAP is required for the activation of both SREBP-1 and SREBP-2 transcription factors (7). Next, we tested whether the requirement of *SCAP* for PDAC cell and tumor growth was due to loss of SREBP-1, SREBP-2, or both. We created *SREBF1* KO*, SREBF2* KO, and *SREBF1/2* double KO Pa03c cell lines and verified these by sequencing and immunoblot analysis (**Fig. 7A**). Deletion of either *SREBF1* or *SREBF2* significantly reduced growth in 1% FBS medium compared to wildtype Pa03c cells **(Fig. 7B)**. Cells lacking both *SREBF1* and *SREBF2* (*BF1/BF2* dKO) showed a greater growth reduction in 1% FBS compared to either *SREBF* single mutant and behaved like *MBTPS1* KO cells that lack the Site-1 protease and all SREBP activity.

**FIGURE 7.**
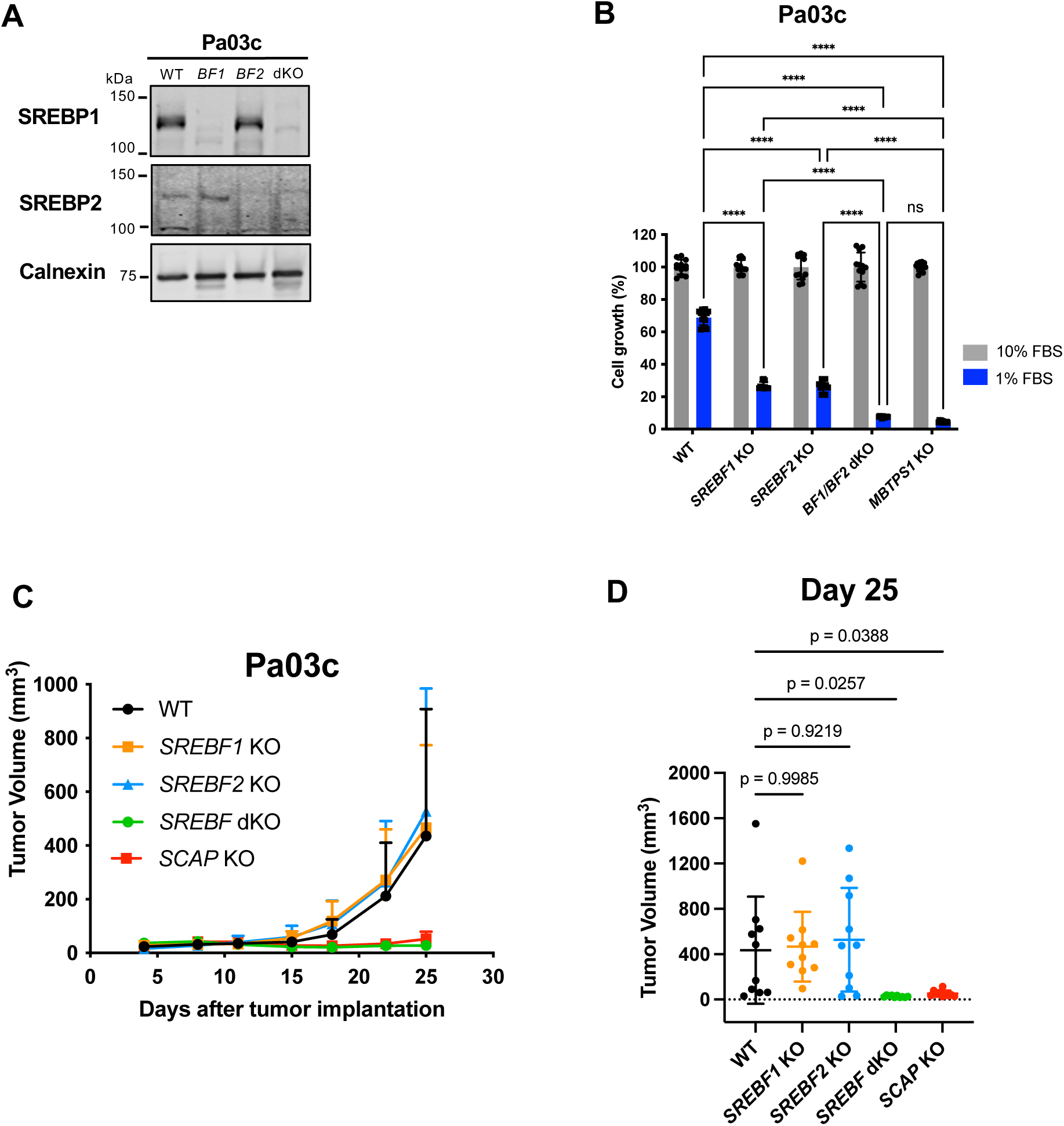
Inhibition of both SREBP-1 and SREBP-2 is required to prevent PDAC cell and tumor growth. **A)** Immunoblot of Pa03c wildtype, *SREBF1* KO, *SREBF2* KO, *SREBF1/2* double KO cells for indicated antibodies. Membrane-enriched extracts (20 µg) were harvested and probed for SREBP1 and SREBP2. Calnexin served as a loading control. **B)** Wildtype Pa03c, *SREBF1* KO*, SREBF2* KO, *SREBF1/2* double KO, and *MBTPS1* KO cells were cultured in either 10% FBS or 1% FBS for 7 days. Media was changed every 3 days. Plates were stained with crystal violet and quantification is shown (n = 3 per group). For each cell line, growth was normalized to the 10% FBS condition. Statistical significance was determined using two-way ANOVA and Tukey’s HSD test. P values are indicated: < 0.05 (*); < 0.01 (**); < 0.001 (***); < 0.0001 (****), not significant (ns). Error bar denotes standard deviation. **C)** Nude mice were subcutaneously injected with 1×10^6^ Pa03c cells of the indicted genotype in both flanks (two tumors per mouse). Once visible, tumors were measured, and volume calculated. Each group contained 5 mice. Error bar denotes standard deviation. **D)** Individual tumor volumes at day 25, n=10 tumors per group. Statistical significance was determined using one-way ANOVA and Dunnett’s test. Error bar denotes standard deviation.

To test the requirement of individual SREBPs for PDAC tumor growth, we assayed growth of single and double KO cells using subcutaneous xenografts in nude mice. Wildtype Pa03c, *SREBF1* KO, and *SREBF2* KO xenografts grew equally well in nude mice (**Fig. 7C-D**). Consistent with our cultured cell experiments (**Fig. 7B**), *SREBF1/2* double KO xenografts phenocopied *SCAP* KO xenografts and failed to grow in nude mice, indicating that the presence of either SREBP-1 or SREBP-2 activity is sufficient to support Pa03c xenograft growth and suggesting that inhibition of both transcription factors is required to prevent PDAC tumor growth. Collectively, these results demonstrate that both *SREBF1* and *SREBF2* support PDAC cell and tumor growth, suggesting that SCAP may be a better therapeutic target than either SREBP-1 or SREBP-2 alone.

### SREBP pathway is broadly required for cancer cell growth and is activated in human PDAC tumors

To examine the requirement of the SREBP pathway for cancer cell growth on a wider scale, we examined the Chronos dependency scores for *SCAP*, *SREBF1*, and *SREBF2* using the Cancer Dependency Map database, which contains genome-wide CRISPR loss-of-function screen data for hundreds of cancer cell lines. A lower dependency score indicates that the gene is more likely to be essential for cell growth. For reference, –1.0 is the median dependency score for all common essential genes. Examining dependency scores for *SCAP,* we found that pancreatic cancer cell lines had the second lowest median dependency score (−1.49), with colorectal cancer cell lines showing the lowest score (−1.51) (**Fig. 8A**). Notably, the median dependency score for *SCAP* was ≤ −1.0 in 7 out of 11 cancer cell types: lung, gastric, breast, urinary tract, prostate, pancreas, and colorectal cancer. Consistent with our cultured cell and subcutaneous xenograft results, neither *SREBF1* nor *SREBF2* individually showed a strong dependency in a parallel analysis (**Fig. 8B-C**). The lowest mean/median dependency scores for *SREBF1* and *SREBF2* in any cancer cell type were −0.5 (pancreas) and −0.2 (prostate), respectively. Therefore, given these data, we conclude that *SCAP* is an essential gene in multiple types of cancers and is likely to be required broadly for growth in pancreatic cancer cell lines.

**FIGURE 8.**
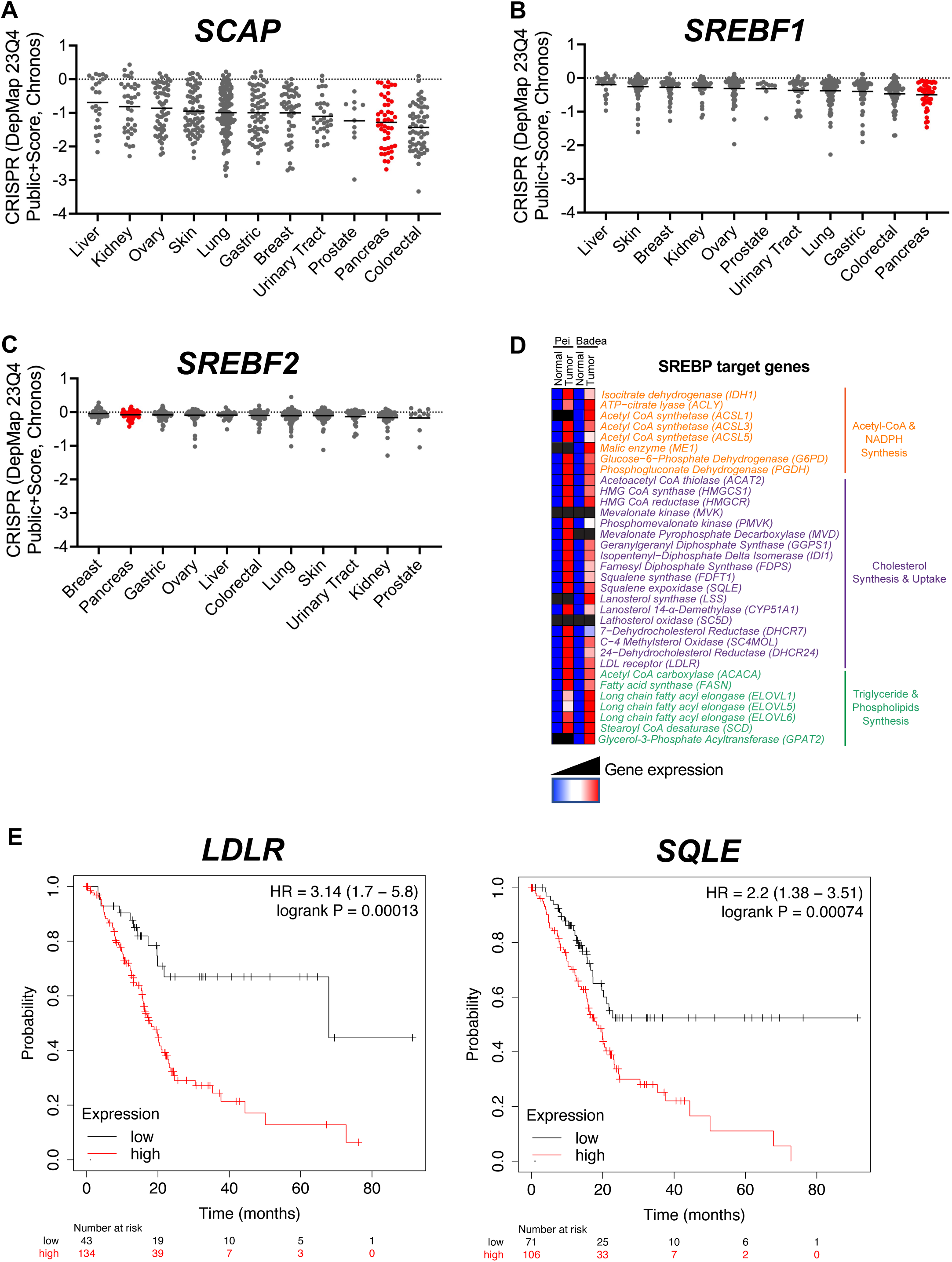
SREBP pathway is broadly required for cancer cell growth and is activated in human PDAC tumors. **A-C)** The essentiality of *SCAP, SREBF1,* and *SREBF2* across cancer cell lines was examined using the Cancer Dependency Map project database (Public Chronos 23Q4). The mean Chronos dependency scores are shown for multiple organ systems, including the pancreas (red). Negative scores indicate gene essentiality. **D)** Expression of SREBP target genes in PDAC tumor tissue compared to normal tissue is shown from two Oncomine data sets Pei and Badea (33). Blue indicates lower gene expression; red indicates higher gene expression; black indicates p> 0.05. Genes are grouped and color-coded by bioinformatic process (https://www.genepattern.org/#gsc.tab=0). **E)** The association of SREBP target genes with PDAC patient survival was queried using the Kaplan-Meier Plotter tool (https://kmplot.com/analysis/) and RNA-seq datasets (n = 177 patients for each curve). HR denotes the hazard ratio.

Considering that the SREBP pathway is required for PDAC xenograft tumor growth, disease progression in KPC mice, and human pancreatic cancer cell lines are highly dependent on *SCAP* function, we next tested whether our observations are applicable to PDAC patients. We compared expression levels of SREBP target genes in PDAC tumors and normal tissue using the Oncomine database (33). In two independent pancreatic cancer data sets, we observed that SREBP target genes were overexpressed in PDAC tumors compared to matched normal tissue, suggesting that pancreatic tumors activate SREBP to support tumor growth (**Fig. 8D**). In addition, we analyzed the association between the expression levels of SREBP target genes and overall survival of PDAC patients (n = 177) using the Kaplan-Meier Plotter Database (34) (**Table S3**). Two SREBP target genes, *low-density lipoprotein receptor (LDLR)* and *squalene epoxidase* (*SQLE*), had an FDR ≤ 5%, and high expression of each gene was significantly associated with worse overall survival in pancreatic cancer patients (p = 0.00013 and p = 0.00074, respectively) (**Fig. 8E**). An independent study of 60 PDAC tumors found that SREBP1 protein expression was elevated in tumor versus normal pancreas tissue and that increased SREBP1 expression correlated with poorer survival (35). Collectively, these observations suggest that SREBP activity is elevated in PDAC tumors compared to normal pancreas tissue and that increased activity may be associated with disease progression and reduced survival.

## DISCUSSION

In this study, we assessed the requirement of the SREBP pathway and its central regulator SCAP in pancreatic ductal adenocarcinoma both *in vivo* and *in vitro*. Using mouse models, human PDAC cell lines and bioinformatics, we demonstrated that 1) loss of *Scap* delays progression to PDAC and prolongs survival in a GEMM of pancreatic cancer (**Fig. 1**), 2) *SCAP* is required for human PDAC subcutaneous and orthotopic tumor growth *in vivo* (**Fig. 2-3**), 3) the SREBP pathway is activated in low serum conditions in PDAC cells (**Fig. 4-6**), 4) the SREBP pathway and *SCAP* are required for PDAC cell growth *in vitro* under low serum and low lipid conditions (**Fig. 4-6**), 5) inhibition of either SREBP-1 or SREBP-2 is not sufficient to inhibit PDAC subcutaneous tumor growth (**Fig. 7**), 6) the SREBP pathway is broadly required for cancer cell growth (**Fig. 8**), and 8) SREBP target genes are upregulated in human PDAC tumors and target gene expression is associated with reduced PDAC patient survival (**Fig. 8**). Taken together, these data indicate that the SREBP pathway and SCAP are potential therapeutic targets for PDAC.

Our results are consistent with parallel studies that tested the requirement for the SREBP pathway both directly and indirectly. Sun and colleagues demonstrated that SREBP-1 protein is elevated in human pancreatic tumors and that this correlated with poor prognosis (35). Knockdown of *SREBF1* slowed growth of human PDAC cells in culture and in subcutaneous xenografts, however, targeting *SREBF1* resulted in incomplete inhibition of cell and tumor growth. Two CRISPR screens for metabolic genes required for mouse PDAC cell growth *in vitro* and *in vivo* also revealed a requirement for *Scap* (36,37). Interestingly, in their screen, Zhu and colleagues found a greater growth dependence *in vivo* for *Scap* than either *Srebf1* or *Srebf2* alone (36). This finding is consistent with our data demonstrating that targeting either *SREBF1* or *SREBF2* is insufficient to prevent human PDAC cell and subcutaneous tumor growth (**Fig. 7**). Inhibiting SCAP instead resulted in complete loss of SREBP pathway activity and more complete growth suppression. One possible explanation for our observation is that cells upregulate activity of the remaining SREBP to compensate, suggesting that SCAP is the preferred therapeutic target.

Xenograft models offer technical ease and the ability to genetically manipulate human PDAC cells that are implanted. However, these models employ nude mice that lack an intact immune system as well as the ability to assess development and progression of tumorigenesis. Using the well-established KPC model, we found that KPC mice had a median survival time of 315 days (**Fig. 1**), in contrast to 172 days in the original model (28). This difference is likely due to the change in background strain from mixed C57Bl/6;129/SvJae to a congenic C57Bl/6 background as well as the application of censoring in the survival curve analysis. Excitingly, mice heterozygous for *Scap* knockout showed a significantly prolonged survival with fewer mice reaching the adverse event than KPC mice (**Fig. 1**). A striking result was noted in the incidence of PDAC and PanINs in these cohorts. While all KPC mice developed the precursor PanIN lesions, none of the homozygous *Scap* knockout mice developed invasive PDAC.

The requirement for the SREBP pathway and Scap for development and organ function has been examined in multiple mouse tissues (38). Broadly, pathway activity is essential for embryonic development as well as maintenance of tissues with high rates of cell division, such as the intestinal epithelium and activated immune cells. Interestingly, Scap is not required for development of mouse liver or mammary epithelium, indicating that basal levels of SREBP target genes are sufficient for development of these tissues (39,40). We developed a pancreas-specific *Scap* knockout mouse to study the role of Scap in PDAC and evaluated *Pdx1-Cre^+/-^ Scap^fl/fl^* mice for effects on pancreas development and function. Our results demonstrate that homozygous loss of *Scap* in the pancreas does not alter the exocrine or endocrine functions of the pancreas (**Supplementary Fig. 1**). Additionally, pancreas development appeared normal. Given the premature death phenotype observed in KPCS^fl/fl^ mice, it is possible that stressing *Pdx1-Cre^+/-^ Scap^fl/fl^* mice would reveal additional phenotypes. However, the fact that *Scap* is not required for normal pancreas development or function provides further evidence that the effects of Scap inhibition are selective for rapidly dividing cells.

It is well established that metabolic reprogramming and upregulation of lipid metabolism is a hallmark of cancer (4). Nutrient-poor tumors upregulate SREBP pathway activity to increase lipid supply and restore lipid homeostasis (**Fig. 8**). Gene expression data from patient PDAC tumors demonstrated that SREBP pathway target genes are upregulated in PDAC tumors compared to normal tissue and that overall survival is reduced in patients whose tumors have high expression of the SREBP target genes *LDLR* and *SLQE* (**Fig. 8**). Consistent with this, Sun and colleagues examined 60 PDAC patient tumors and found increased expression of SREBP-1 in tumors versus normal tissue, confirming our observations (14). In our studies, chemical and genetic inhibition of the SREBP pathway in human PDAC cells reduced gene expression of SREBP target genes in fatty acid synthesis, the mevalonate pathway, and cholesterol uptake and inhibited PDAC cell growth (**Fig. 4-6**). Given that SREBPs activate a broad program of lipid metabolism gene expression (6), inhibition of this adaptive response likely results in disruption of lipid homeostasis, preventing PDAC cell and tumor growth.

Lipid synthesis enzymes have been explored as cancer therapeutic targets for several decades (41,42). These approaches seek to eliminate the activity of a single enzyme or metabolic pathway. Targeting SCAP represents a different approach, as loss of SREBP activity does not alter basal expression of target genes. Rather, inhibition of SREBPs prevents the upregulation of lipid supply pathways under nutrient-poor conditions and subsequent metabolic adaptation needed to support tumor growth. Consistent with the requirement of *SCAP* for PDAC tumor growth, multiple SREBP target genes have individually been shown to be required for PDAC tumor growth: *ACSL3*, *ACLY*, *FDFT1*, *GGPS1*, NSDHL, and *LDLR* (37,40,43–46). It remains to be determined whether the observed effects of SCAP inhibition on PDAC are the result of failure to upregulate a single gene and pathway product or whether this represents the combined effects of multiple deficiencies. Understanding the mechanism underlying the requirement for SCAP may reveal vulnerabilities that present opportunities for combination therapy.

The SREBP pathway has a demonstrated requirement in a growing list of cancers such as colon, gastric, breast, prostate, and glioblastoma (13–17). Consistent with this, we find a strong growth dependence on SCAP in multiple cancers (**Fig. 8A**). Our results indicate that PDAC should be added to this list. Here, we demonstrate a requirement for SCAP in the development and progression of PDAC under conditions when SCAP function is removed during carcinogenesis. It remains to be tested whether inhibition of SCAP in established tumors prevents growth, metastasis, and prolongs survival. As such, future studies should employ both genetic and pharmacological inhibition of the SREBP pathway in established primary and metastatic PDAC tumors. Given the requirement for the SREBP pathway in a wide range of solid tumors, development of a potent, bioavailable chemical inhibitor is critical and will enable evaluation of the SREBP pathway as a therapeutic target in preclinical cancer models.

## AUTHORS’ CONTRIBUTIONS

**C. Ishida**: Conceptualization, formal analysis, investigation, visualization, writing-original draft. **S.L. Myers**: Conceptualization, formal analysis, investigation, visualization, writing-original draft, funding acquisition. **W. Shao**: Conceptualization, formal analysis, investigation, visualization. **M.R. McGuire**: Conceptualization, formal analysis, investigation, writing-review and editing. **C. Liu**: Conceptualization, formal analysis, investigation, visualization. **C.S. Kubota**: Formal analysis, investigation, visualization, writing-review and editing. **T.E. Ewachiw**: Investigation. **D. Mukhopadhyay**: Formal analysis, writing-review and editing. **S. Ke**: Formal analysis, visualization. **H. Wang**: Formal analysis, supervision. **Z.A. Rasheed**: Supervision. **R.A. Anders**: Investigation. **P.J. Espenshade**: Conceptualization, funding acquisition, project administration, supervision, writing-original draft.

## ACKNOWLEDGEMENTS

We thank Anirban Maitra (Department of Pathology, Johns Hopkins School of Medicine; current appointment, MD Anderson Cancer Center) for providing human PDAC cell lines; William Matsui (Department of Oncology, Johns Hopkins School of Medicine; current appointment, Department of Oncology, Dell Medical School, University of Texas) for experimental advice; and Lei Zheng (Department of Oncology, Johns Hopkins University School of Medicine) for providing the KPC breeder mice.

## REFERENCES

1. Siegel RL, Miller KD, Wagle NS, Jemal A. Cancer statistics, 2023. CA: A Cancer Journal for Clinicians 2023;73:17-48.

2. Abbassi R, Algül H. Palliative chemotherapy in pancreatic cancer-treatment sequences. Translational Gastroenterology and Hepatology 2019;4:6–11.

3. Hosein AN, Dougan SK, Aguirre AJ, Maitra A. Translational advances in pancreatic ductal adenocarcinoma therapy. Nature Cancer 2022;3:272–86.

4. Ward PS, Thompson CB. Metabolic reprogramming: a cancer hallmark even Warburg did not anticipate. Cancer Cell 2012;21:297–308.

5. Snaebjornsson MT, Janaki-Raman S, Schulze A. Greasing the wheels of the cancer machine: The role of lipid metabolism in cancer. Cell Metab 2020;31:62–76.

6. Horton JD, Shah NA, Warrington JA, Anderson NN, Park SW, Brown MS, et al. Combined analysis of oligonucleotide microarray data from transgenic and knockout mice identifies direct SREBP target genes. Proceedings of the National Academy of Sciences of the United States of America 2003;100:12027–32.

7. Shimano H, Sato R. SREBP-regulated lipid metabolism: convergent physiology - divergent pathophysiology. Nat Rev Endocrinol 2017;13:710–30.

8. Ricoult SJ, Yecies JL, Ben-Sahra I, Manning BD. Oncogenic PI3K and K-Ras stimulate de novo lipid synthesis through mTORC1 and SREBP. Oncogene 2016:1250–60.

9. Ying H, Dey P, Yao W, Kimmelman AC, Draetta GF, Maitra A, et al. Genetics and biology of pancreatic ductal adenocarcinoma. Genes Dev 2016;30:355–85.

10. Hruban RH, Adsay NV, Albores-Saavedra J, Anver MR, Biankin AV, Boivin GP, et al. Pathology of genetically engineered mouse models of pancreatic exocrine cancer: consensus report and recommendations. Cancer Research 2006;66:95–106.

11. Sodir NM, Kortlever RM, Barthet VJA, Campos T, Pellegrinet L, Kupczak S, et al. MYC instructs and maintains pancreatic adenocarcinoma phenotype. Cancer discovery 2020;10:588–607.

12. Gouw AM, Margulis K, Liu NS, Raman SJ, Mancuso A, Toal GG, et al. The MYC oncogene cooperates with Sterol-Regulated Element-Binding Protein to regulate lipogenesis essential for neoplastic growth. Cell Metab 2019;30:556–72 e5.

13. Wen YA, Xiong XP, Zaytseva YY, Napier DL, Vallee E, Li AT, et al. Downregulation of SREBP inhibits tumor growth and initiation by altering cellular metabolism in colon cancer. Cell Death Dis 2018;9:265.

14. Sun Q, Yu X, Peng C, Liu N, Chen W, Xu H, et al. Activation of SREBP-1c alters lipogenesis and promotes tumor growth and metastasis in gastric cancer. Biomedicine & Pharmacotherapy 2020;128:110274.

15. Bao JS, Zhu LP, Zhu Q, Su JH, Liu ML, Huang W. SREBP-1 is an independent prognostic marker and promotes invasion and migration in breast cancer. Oncol Lett 2016;12:2409–16.

16. Krycer JR, Phan L, Brown AJ. A key regulator of cholesterol homoeostasis, SREBP-2, can be targeted in prostate cancer cells with natural products. Biochem J 2012;446:191–201.

17. Lewis CA, Brault C, Peck B, Bensaad K, Griffiths B, Mitter R, et al. SREBP maintains lipid biosynthesis and viability of cancer cells under lipid- and oxygen-deprived conditions and defines a gene signature associated with poor survival in glioblastoma multiforme. Oncogene 2015;34:5128–40.

18. Cheng C, Geng F, Cheng X, Guo D. Lipid metabolism reprogramming and its potential targets in cancer. Cancer Communications 2018 38:1 2018;38:1–14.

19. Feldmann G, Beaty R, Hruban RH, Maitra A. Molecular genetics of pancreatic intraepithelial neoplasia. Journal of Hepato-Biliary-Pancreatic Surgery 2007;14:224–32.

20. Jones S, Zhang X, Parsons DW, Lin JC, Leary RJ, Angenendt P, et al. Core signaling pathways in human pancreatic cancers revealed by global genomic analyses. Science 2008;321:1801–6.

21. Hingorani SR, Petricoin EF, Maitra A, Rajapakse V, King C, Jacobetz MA, et al. Preinvasive and invasive ductal pancreatic cancer and its early detection in the mouse. Cancer Cell 2003;4:437–50.

22. Matsuda M, Korn BS, Hammer RE, Moon YA, Komuro R, Horton JD, et al. SREBP cleavage-activating protein (SCAP) is required for increased lipid synthesis in liver induced by cholesterol deprivation and insulin elevation. Genes Dev 2001;15:1206–16.

23. Foley K, Rucki AA, Xiao Q, Zhou D, Leubner A, Mo G, et al. Semaphorin 3D autocrine signaling mediates the metastatic role of annexin A2 in pancreatic cancer. Sci Signal 2015;8:ra77.

24. Tomayko MM, Reynolds CP. Determination of subcutaneous tumor size in athymic (nude) mice. Cancer Chemother Pharmacol 1989;24:148–54.

25. Treuting PM, Dintzis SM, Montine KS. Comparative Anatomy And Histology: A Mouse, Rat, And Human Atlas Second Edition. 2018:543.

26. Fay MP, Shaw PA. Exact and asymptotic weighted logrank tests for interval censored data: The interval R package. J Stat Softw 2010;36

27. Carr RM, Fernandez-Zapico ME. Pancreatic cancer microenvironment, to target or not to target? Embo Mol Med 2016;8:80–2.

28. Hingorani SR, Wang L, Multani AS, Combs C, Deramaudt TB, Hruban RH, et al. Trp53R172H and KrasG12D cooperate to promote chromosomal instability and widely metastatic pancreatic ductal adenocarcinoma in mice. Cancer Cell 2005;7:469–83.

29. Hu HF, Ye Z, Qin Y, Xu XW, Yu XJ, Zhuo QF, et al. Mutations in key driver genes of pancreatic cancer: molecularly targeted therapies and other clinical implications. Acta Pharmacologica Sinica 2021;42:1725–41.

30. Hawkins JL, Robbins MD, Warren LC, Xia D, Petras SF, Valentine JJ, et al. Pharmacologic inhibition of site 1 protease activity inhibits sterol regulatory element-binding protein processing and reduces lipogenic enzyme gene expression and lipid synthesis in cultured cells and experimental animals. J Pharmacol Exp Ther 2008;326:801–8.

31. Radhakrishnan A, Ikeda Y, Kwon HJ, Brown MS, Goldstein JL. Sterol-regulated transport of SREBPs from endoplasmic reticulum to Golgi: Oxysterols block transport by binding to Insig. Proceedings of the National Academy of Sciences of the United States of America 2007;104:6511–8.

32. Janowski BA, Grogan MJ, Jones SA, Wisely GB, Kliewer SA, Corey EJ, et al. Structural requirements of ligands for the oxysterol liver X receptors LXRa and LXRβ. Proceedings of the National Academy of Sciences of the United States of America 1999;96:266–71.

33. Rhodes DR, Yu JJ, Shanker K, Deshpande N, Varambally R, Ghosh D, et al. ONCOMINE: A cancer microarray database and integrated data-mining platform. Neoplasia 2004;6:1–6.

34. Lánczky A, Győrffy B. Web-based survival analysis tool tailored for medical research (KMplot): development and implementation. Journal of Medical Internet Research 2021;23:e27633.

35. Sun Y, He W, Luo M, Zhou Y, Chang G, Ren W, et al. SREBP1 regulates tumorigenesis and prognosis of pancreatic cancer through targeting lipid metabolism. Tumor Biology 2015;36:4133–41.

36. Zhu XG, Chudnovskiy A, Baudrier L, Prizer B, Liu Y, Ostendorf BN, et al. Functional genomics in vivo reveal metabolic dependencies of pancreatic cancer cells. Cell Metabolism 2021;33:211–21.

37. Biancur DE, Kapner KS, Yamamoto K, Banh RS, Neggers JE, Sohn ASW, et al. Functional genomics identifies metabolic vulnerabilities in pancreatic cancer. Cell Metabolism 2021;33:199–210.

38. Engelking LJ, Cantoria MJ, Xu YC, Liang GS. Developmental and extrahepatic physiological functions of SREBP pathway genes in mice. Seminars in Cell & Developmental Biology 2018;81:98–109.

39. Moon YA, Liang G, Xie X, Frank-Kamenetsky M, Fitzgerald K, Koteliansky V, et al. The Scap/SREBP pathway is essential for developing diabetic fatty liver and carbohydrate-induced hypertriglyceridemia in animals. Cell Metab 2012;15:240–6.

40. Sebastiano MR, Pozzato C, Saliakoura M, Yang Z, Peng RW, Galiè M, et al. ACSL3-PAI-1 signaling axis mediates tumor-stroma cross-talk promoting pancreatic cancer progression. Science advances 2020;6

41. Menendez JA, Lupu R. Fatty acid synthase and the lipogenic phenotype in cancer pathogenesis. Nat Rev Cancer 2007;7:763–77.

42. Mullen PJ, Yu R, Longo J, Archer MC, Penn LZ. The interplay between cell signalling and the mevalonate pathway in cancer. Nat Rev Cancer 2016;16:718–31.

43. Carrer A, Trefely S, Zhao S, Campbell SL, Norgard RJ, Schultz KC, et al. Acetyl-CoA metabolism supports multistep pancreatic tumorigenesis. Cancer discovery 2019;9:416–35.

44. Guillaumond F, Bidaut G, Ouaissi M, Servais S, Gouirand V, Olivares O, et al. Cholesterol uptake disruption, in association with chemotherapy, is a promising combined metabolic therapy for pancreatic adenocarcinoma. Proceedings of the National Academy of Sciences of the United States of America 2015;112:2473–8.

45. Haney SL, Varney ML, Chhonker YS, Shin S, Mehla K, Crawford AJ, et al. Inhibition of geranylgeranyl diphosphate synthase is a novel therapeutic strategy for pancreatic ductal adenocarcinoma. Oncogene 2019 38:26 2019;38:5308-20.

46. Gabitova-Cornell L, Surumbayeva A, Peri S, Franco-Barraza J, Restifo D, Weitz N, et al. Cholesterol pathway inhibition induces TGF-β signaling to promote basal differentiation in pancreatic cancer. Cancer Cell 2020;38:567–83.e11.

