## Supplemental Methods and Figures for "SREBP-dependent regulation of lipid homeostasis is required for progression and growth of pancreatic ductal adenocarcinoma"

#### SUPPLEMENTARY MATERIALS AND METHODS

##### Chemical Reagents

We obtained chemicals from the following manufacturers with catalog numbers in parenthesis: PF-429242 dihydrochloride (MedChemExpress, HY-13447A), 25-hydroxycholesterol (MilliporeSigma, H1015), cholesterol (MilliporeSigma, C3045), LDL (Prospec Bio, PRO-562), puromycin (MilliporeSigma, P8833), blasticidin (Corning, 30100RB), crystal violet (MilliporeSigma, C3886). Other general chemicals were obtained from MilliporeSigma or Thermo Fisher Scientific.

##### Generation of CRISPR/Cas9-mediated Knockout Cells

*SCAP* KO Pa02c, Pa03c, Pa16c, and Pa20c cells were generated using CRISPR/Cas9 genome editing (1). *SCAP* KO Pa03c cells were generated as previously described using blasticidin selection and the guide sequence below (2). For the generation of other KO cells, guide RNA sequence targeting *SCAP* (5'-GGCTGCGTGAGAAGATATCT -3'), *SREBF1* (5'- GGAGGTGGAGACAAGCTGCC -3'), *SREBF2* (5'- GCAGACCCTTGCCCCGGCTA -3') or *MBTPS1* (5'-GTGCGGGGTTCTGGCGTGAA -3') was cloned into the Cas9-gRNA vector PX459 (Addgene 48139). Cell lines names are Pa03c SCAP KO (WSC55), Pa03c BP1 KO (WSC139), Pa03c BP2 KO (WSC142), Pa03c BP1/BP2 DKO (WSC120), Pa03c S1P KO (WSC104), Pa16c SCAP KO (WSC143), Pa02c SCAP KO (WSC145), and Pa20c SCAP KO (WSC146). Transduced Pa cells were selected in M19 medium containing 1.5 µg/mL puromycin until the control cells without transfection died. Single clones were isolated by dilution cloning. Genomic DNA flanking the gRNA target site was amplified by standard PCR and sequenced by Sanger sequencing. Knockout of *SCAP* was confirmed by immunoblotting and growth assays under lipoprotein-depleted conditions. For the generation of *SCAP* rescue cells, guide RNA-resistant human *SCAP* sequence (c.36T>C, c.39G>C) was cloned into the following plasmids: Cas9 plasmid (Addgene 52962 for Pa02c, Pa16c, and Pa20c cells) after removing Cas9 by restriction enzyme digestion and lentiGuide-Puro plasmid Addgene 52963 for Pa03c cells. Transduced *SCAP* KO cells were selected in DMEM medium supplemented with 10% FBS (v/v) containing blasticidin (Pa02c, Pa16c, and Pa20c cells) or puromycin (Pa03c cells). SCAP expression was confirmed by immunoblotting and growth assays under lipoprotein-depleted conditions.

#### Real-Time qPCR

Total RNA was extracted using Monarch$^{R}$ Total RNA Miniprep Kit (New England BioLabs, T2010S) and reverse transcription was carried out using iScript$^{TM}$ Reverse Transcription Supermix (Bio-Rad, 1708841) according to the manufacturer's protocol. Target gene expression was evaluated by real-time qPCR using SYBER Green Master Mix (Thermofisher, A25778) according to manufacturer's protocol with primers listed in **Table S4**. *GAPDH* was used as the internal control to calculate the relative expression of target genes from different samples.

#### Crystal Violet Growth Assay

Crystal violet growth assay has been previously described (3). Briefly, $5x10^3$ cells were seeded in 24-well plates 24 hours before indicated treatment. Medium was replaced every 48 hours. After the indicated experimental time, medium was aspirated, and cells were washed with ice-cold PBS. Cells were then fixed with ice cold methanol (stored at -20°C) and incubated for 10 minutes at -20°C. The methanol was aspirated, and 0.05% (w/v) crystal violet solution in 25% (v/v) methanol was added and allowed to stain for 10 minutes at room temperature. The cells were gently washed with deionized water and plates were dried overnight at room temperature. Stained plates were scanned using an Epson scanner to obtain images. For quantification, 10% (v/v) fresh acetic acid was added to each well, and the plate was incubated for 15 minutes at room temperature. Acetic acid-dye solution (100 µL) was transferred to a 96-well plate, and absorbance was measured at 590 nm using FLUOstar Omega microplate reader (BMG LABTECH).

**Cell Proliferation Assay**

Cells were seeded at $3x10^3$ in 100 µL of medium in 96-well plates 24 hours before indicated treatment. Cells were then grown for 72 hours. To assay viable cells, 20 µL of MTS reagent (CellTiter 96 AQueous One Solution Proliferation Assay, Promega, G358Taq) was added to each well, and plate was incubated at 37˚C for 3.5 hours. Absorbance was measured at 490 nm using FLUOstar Omega microplate reader (BMG LABTECH).

**Mouse Husbandry and Genotyping**

Routine health surveillance using dirty bedding sentinel serology indicated that the mice were free of the following organisms: mouse hepatitis virus, minute virus of mice, mouse parvovirus, epizootic diarrhea of infant mice (rotavirus), Theiler's murine encephalomyelitis virus, murine norovirus, Sendai virus, pneumonia virus of mice, reovirus, lymphocytic choriomeningitis virus, ectromelia virus, mouse adenovirus (FL & K87), mouse cytomegalovirus, *Mycoplasma pulmonis*, fur mites, and pinworms. Mice were housed in social groups (2-5 mice) of the same sex in individually ventilated cages (Allentown Caging Inc., Allentown, NJ) with autoclaved corncob bedding (Teklad, Envigo, NJ) and nesting material (Animal Specialties and Provisions, Quakertown, PA). Mice are visually assessed every morning, and cages were changed every 14 days. Autoclaved feed (Teklad Global 2018S) was provided *ad libitum*, and water was provided via an in-cage automated watering system (Systems Engineering, Inc., Napa, CA). The room was maintained at 22±1°C on a 14:10 light:dark cycle at 40-70% humidity. Unless otherwise stated, euthanasia was performed using either carbon dioxide asphyxiation at 30-70% displacement or isoflurane anesthesia followed by either cervical dislocation or exsanguination and bilateral thoracotomy (4).

To obtain maximal breeding efficiency and avoid perineal masses that developed in KC females, the following crosses were used: for KPC (*Kras*$^{LSL\text{-}G12D/+}$;*Trp53*$^{LSL\text{-}R172H/+}$;*Pdx1-Cre*$^{+/-}$) mice, KC males were bred to KP females; for KPCS$^{fl/+}$ (*Kras*$^{LSL\text{-}G12D/+}$;*Trp53*$^{LSL\text{-}R172H/+}$;*Pdx1-Cre*$^{+/-}$; *Scap*$^{fl/+}$) mice, KCS$^{fl/+}$ males were crossed to KPS$^{fl/+}$ females; for KPCS$^{fl/fl}$ (*Kras*$^{LSL\text{-}G12D/+}$;*Trp53*$^{LSL\text{-}R172H/+}$;*Pdx1-Cre*$^{+/-}$; *Scap*$^{fl/fl}$) mice, KCS$^{fl/fl}$ males were crossed to KPS$^{fl/fl}$ females.

Multiplex genotyping of non-pancreatic [tail biopsy (<5 mm) or ear punch (2 mm)] was performed at weaning. DNA was isolated using alkaline hydrolysis. Briefly, tissue was incubated in 50 µL lysis solution (25 mM NaOH and 0.2 mM EDTA) at 90ºC for 10-60 minutes, then neutralized with 50 µL 40 mM Tris-HCl (pH 5.5). Genotyping PCR was performed using indicated primers (**Table S5**), GoTaq Hot Start Master Mix (Promega, M5123) using the manufacturer's protocol , and 1 µL of isolated DNA with the following PCR cycle conditions: 94°C for 3 minutes; 38 cycles of denaturing at 94°C for 30 seconds, annealing at 60°C for 30 seconds, elongation at 72°C for 25 seconds; final elongation at 72°C for 30 seconds; cooled to 4°C. Genotyping of pancreatic tissue was performed using the indicated primers (**Table S5**), 5X Green GoTaq Buffer (Promega, M7911) and up to 5 units of Choice Taq polymerase (Denville, CB4050-1) using the above PCR cycle conditions with the following modifications: annealing at 55°C, 35 cycles. PCR products were run on a 2% agarose gel for band identification.

**Glucose and Insulin Tolerance Tests**

Male and female C, $CS^{fl/fl}$, $S^{fl/fl}$, or wildtype C57BL/6J mice aged 6 weeks old were bred and maintained on standard rodent chow (Teklad Global, 2018S). Intraperitoneal glucose and insulin tolerance tests were performed as previously described (5). Briefly, mice were fasted, but provided free access to water, from 9 am to 3 pm (total of 6 hours). Baseline blood glucose levels were measured in tail vein blood using a Bayer Contour glucometer. Mice were then injected with 2 mg/kg of D-Glucose (Fisher Scientific, D16-10) or 0.5 IU/kg of Humulin R Insulin (Eli Lilly, U100 or NDC 0002-8215-01) intraperitoneally for glucose tolerance and insulin tolerance tests, respectively. Blood glucose levels were then re-measured at 15, 30, 60, 90, and 120 minutes post-injection.

**Tissue Membrane Fractionation**

Tissue membrane fractionation was performed as previously described (6). Briefly, livers and pancreases were harvested from male C, $CS^{fl/fl}$, $S^{fl/fl}$, or wildtype C57BL/6J mice aged 4-12 months and frozen by liquid nitrogen. Frozen tissues were homogenized in buffer (20 mM Tris-HCl, pH 7.4, 2 mM $MgCl_2$, 0.25 mM sucrose, 10 mM EDTA, 10 mM EGTA) with 1 mM PMSF and Halt protease and phosphatase inhibitor cocktail (Thermo Fisher Scientific, PI78442). Homogenized tissues were centrifuged at 1 000 x g for 5 minutes at 4°C and the supernatant was centrifuged at 20 000 x g for 1 hour at 4°C. The resulting membrane pellets were resuspended in SDS-lysis buffer (10 mM Tris-HCl, pH 6.8, 100 mM NaCl, 1% (w/v) SDS, 1 mM EDTA, 1 mM EGTA) with 1 mM PMSF and Halt protease and phosphatase inhibitor cocktail (Thermo Fisher Scientific), and the protein concentration was measured using the BCA Protein Assay Kit (Pierce). Membrane fraction was diluted with SDS-lysis buffer and mixed with an equal volume of urea buffer (62.5 mM Tris-HCl, pH 6.8, 15% (w/v) SDS, 8 M urea, 10% (v/v) glycerol, 100 mM DTT). Membrane proteins were separated by SDS-PAGE and transferred to nitrocellulose membranes as described under Immunoblot Analysis.

**Protein Extraction and Immunoblot Analysis**

Cultured cells were homogenized in SDS-lysis buffer (10 mM Tris-HCl, pH 6.8, 100 mM NaCl, 1% (w/v) SDS, 1 mM EDTA, 1 mM EGTA) with protease inhibitors (5 µg/mL pepstatin A, 10 µg/mL leupeptin, 0.5 mM PMSF, 1 mM DTT) for generating whole cell protein extracts. Cell fractionation (membrane and nuclear) was performed as previously described (7). Protein concentration was measured with a BCA Protein Assay Kit (Pierce, 23227). Proteins were separated by SDS-PAGE and transferred to nitrocellulose membranes using the Trans-Blot Turbo Transfer system (Bio-Rad). After blocking in 5% (w/v) instant nonfat dry milk (Nestle) in PBST (1X PBS with 0.5% (v/v) Tween20, MilliporeSigma BP337500) for 30 minutes at room temperature, membranes were incubated with primary antibodies. Antibodies were diluted in PBST and incubated at room temperature for 1 hour or at 4°C overnight. The following antibodies were used: SREBP-1 2A4 (1:40, Santa Cruz Biotechnology, SC-13551), SREBP-2 22D5 (1:10 000) (8), KDM1A (1:1 000, Cell Signaling, 2184), SCD (1:1 000, Abcam, 19862), LDLR (1:500, abcam, 52818), Actin (1:1 000, Santa Cruz Biotechnology, SC-47778), SCAP (1:1 000) (9), Calnexin (1:1 000, Sigma, C7617). Membranes were then washed at least 3 times in PBST for 5 minutes each at room temperature followed by incubation with secondary antibody at room temperature for 30 minutes. Secondary antibodies include IRDye 800CW or IRDye 680RD-conjugated goat anti-mouse or anti-rabbit IgG (1:20 000 in 1% (w/v) non-fat milk in PBST (Li-COR Biosciences, NC9401841, NC9401842). Detection of fluorescence was performed using Odyssey CLx imaging system (LI-COR Biosciences).

**Microarray Analysis**

Pa03c cells were cultured in DMEM medium supplemented with 10% (v/v) FBS or 0.5% (v/v) FBS for 16 hours. RNA samples were harvested using the QIAGEN RNeasy RNA kit (7410). Genome-scale gene expression measurement was conducted by the Sidney Kimmel Comprehensive Cancer Center at the Johns Hopkins Microarray Core Facility using Illumina HumanHT-12 v4 Expression bead arrays. The resultant Illumina idat files were imported in R (v4.1.3), then normalized and background corrected using the read.idat (using HumanHT-12_V4_0_R2_15002873_B.bgx manifest file) and neqc functions as implemented in limma package (v3.46.0) (10). Genes with expression values < 0.01 were removed from the dataset, followed by ranking of the genes based on fold change in expression (0.5% FBS vs 10% FBS). This pre-ranked gene list was used in GSEA (v4.1.0) along with MSigDB (v7.4) Hallmark, Reactome, KEGG and Wikipathways gene sets (11-16).

**Bioinformatic Analysis**

The expression data of SREBP target genes in pancreatic tumor tissue compared to normal tissue were obtained from two Oncomine data sets, Pei (17) and Badea (18,19), and processed using GenePattern (20). The correlation between the expression of SREBP target genes, *LDLR* and *SQLE* (mRNA) and PDAC patient (n=177) survival was assessed using the Kaplan-Meier Plotter tool (21). For gene dependency analysis, data were obtained from the Public Chronos 23Q4 dataset in the Cancer Dependency Map database (22,23), available online at https://depmap.org/portal/download/all/. Gene dependency scores for *SCAP, SREBF1,* and *SREBF2* across 571 cancer cell lines were downloaded and plotted by lineage.

**Photography and Histology**

All gross photographs were taken using either a Sony a6300 digital camera or iPhone 12 Pro. Images were edited for brightness, contrast, and color saturation across the whole image in Adobe Photoshop. Routine processing, paraffin embedding, sectioning (4 µm), and hematoxylin and eosin staining were performed according to standard protocols by either the Department of Molecular and Comparative Pathobiology or Reference Histology Core Facility (Johns Hopkins University). All histologic analysis was performed by a board-certified veterinary pathologist. Slides were viewed on a Nikon Eclipse Ci microscope, and images captured using a Nikon DS-Fi2 camera. Images were edited for white balance across the whole image in Adobe Photoshop.

#### REFERENCES


1. Jinek M, East A, Cheng A, Lin S, Ma E, Doudna J. RNA-programmed genome editing in human cells. eLife **2013**;2:e00471.

2. Bhattacharjee A, Yang H, Duffy M, Robinson E, Conrad-Antoville A, Lu YW*, et al.* The activity of Menkes Disease protein ATP7A is essential for redox balance in mitochondria. J Biol Chem **2016**;291:16644-58.

3. Shao W, Machamer CE, Espenshade PJ. Fatostatin blocks ER exit of SCAP but inhibits cell growth in a SCAP-independent manner. J Lipid Res **2016**;57:1564-73.

4. Leary S, Underwood W, Anthony RA, Cartner S, Grandin T, Greenacre C*, et al.* AVMA Guidelines for the Euthanasia of Animals: 2020 Edition. 2020. 1-121 p.

5. Song WJ, Seshadri M, Ashraf U, Mdluli T, Mondal P, Keil M*, et al.* Snapin mediates incretin action and augments glucose-dependent insulin secretion. Cell Metab **2011**;13:308-19.

6. Engelking LJ, Kuriyama H, Hammer RE, Horton JD, Brown MS, Goldstein JL*, et al.* Overexpression of Insig-1 in the livers of transgenic mice inhibits SREBP processing and reduces insulin-stimulated lipogenesis. JClinInvest **2004**;113:1168-75.

7. DeBose-Boyd RA, Brown MS, Li WP, Nohturfft A, Goldstein JL, Espenshade PJ. Transport-dependent proteolysis of SREBP: Relocation of Site-1 protease from Golgi to ER obviates the need for SREBP transport to Golgi. Cell **1999**;99:703-12.

8. Seo YK, Jeon TI, Chong HK, Biesinger J, Xie X, Osborne TF. Genome-wide localization of SREBP-2 in hepatic chromatin predicts a role in autophagy. Cell Metab **2011**;13:367-75.

9. Sakai J, Nohturfft A, Cheng D, Ho YK, Brown MS, Goldstein JL. Identification of complexes between the COOH-terminal domains of sterol regulatory element-binding proteins (SREBPs) and SREBP cleavage-activating protein. Journal of Biological Chemistry **1997**;272:20213-21.

10. Ritchie ME, Phipson B, Wu D, Hu Y, Law CW, Shi W*, et al.* limma powers differential expression analyses for RNA-sequencing and microarray studies. Nucleic Acids Res **2015**;43:e47.

11. Kanehisa M, Furumichi M, Tanabe M, Sato Y, Morishima K. KEGG: new perspectives on genomes, pathways, diseases and drugs. Nucleic Acids Research **2017**;45:D353-D61.

12. Mootha VK, Lindgren CM, Eriksson KF, Subramanian A, Sihag S, Lehar J*, et al.* PGC-1alpha-responsive genes involved in oxidative phosphorylation are coordinately downregulated in human diabetes. Nat Genet **2003**;34:267-73.

13. Subramanian A, Tamayo P, Mootha VK, Mukherjee S, Ebert BL, Gillette MA*, et al.* Gene set enrichment analysis: a knowledge-based approach for interpreting genome-wide expression profiles. Proc Natl Acad Sci U S A **2005**;102:15545-50.

14. Liberzon A, Birger C, Thorvaldsdóttir H, Ghandi M, Mesirov JP, Tamayo P. The Molecular Signatures Database (MSigDB) hallmark gene set collection. Cell Systems **2015**;1:417-25.

15. Fabregat A, Jupe S, Matthews L, Sidiropoulos K, Gillespie M, Garapati P*, et al.* The Reactome Pathway Knowledgebase. Nucleic Acids Research **2018**;46:D649-D55.

16. Slenter DN, Kutmon M, Hanspers K, Riutta A, Windsor J, Nunes N*, et al.* WikiPathways: a multifaceted pathway database bridging metabolomics to other omics research. Nucleic Acids Research **2018**;46:D661-D7.

17. Pei H, Li L, Fridley BL, Jenkins GD, Kalari KR, Lingle W*, et al.* FKBP51 affects cancer cell response to chemotherapy by negatively regulating Akt. Cancer Cell **2009**;16:259-66.

18. Badea L, Herlea V, Dima S, Dumitrașcu T, Popescu I. Combined gene expression analysis of whole-tissue and microdissected pancreatic ductal adenocarcinoma identifies genes specifically overexpressed in tumor epithelia. Hepato-gastroenterology **2008**;55:2016-27.

19. Rhodes DR, Yu JJ, Shanker K, Deshpande N, Varambally R, Ghosh D*, et al.* ONCOMINE: A cancer microarray database and integrated data-mining platform. Neoplasia **2004**;6:1-6.

20. Reich M, Liefeld T, Gould J, Lerner J, Tamayo P, Mesirov JP. GenePattern 2.0. Nat Genet **2006**;38:500-1.

21. Lánczky A, Győrffy B. Web-based survival analysis tool tailored for medical research (KMplot): development and implementation. Journal of Medical Internet Research **2021**;23:e27633.

22. Dempster JM, Boyle I, Vazquez F, Root DE, Boehm JS, Hahn WC*, et al.* Chronos: a cell population dynamics model of CRISPR experiments that improves inference of gene fitness effects. Genome Biology **2021**;22:1-23.

23. DepMap, Broad. Volume DepMap 23Q4 Public. figshare. Dataset. 2023.

24. Matsuda M, Korn BS, Hammer RE, Moon YA, Komuro R, Horton JD*, et al.* SREBP cleavage-activating protein (SCAP) is required for increased lipid synthesis in liver induced by cholesterol deprivation and insulin elevation. Genes Dev **2001**;15:1206-16.

### Supplementary Figure 1

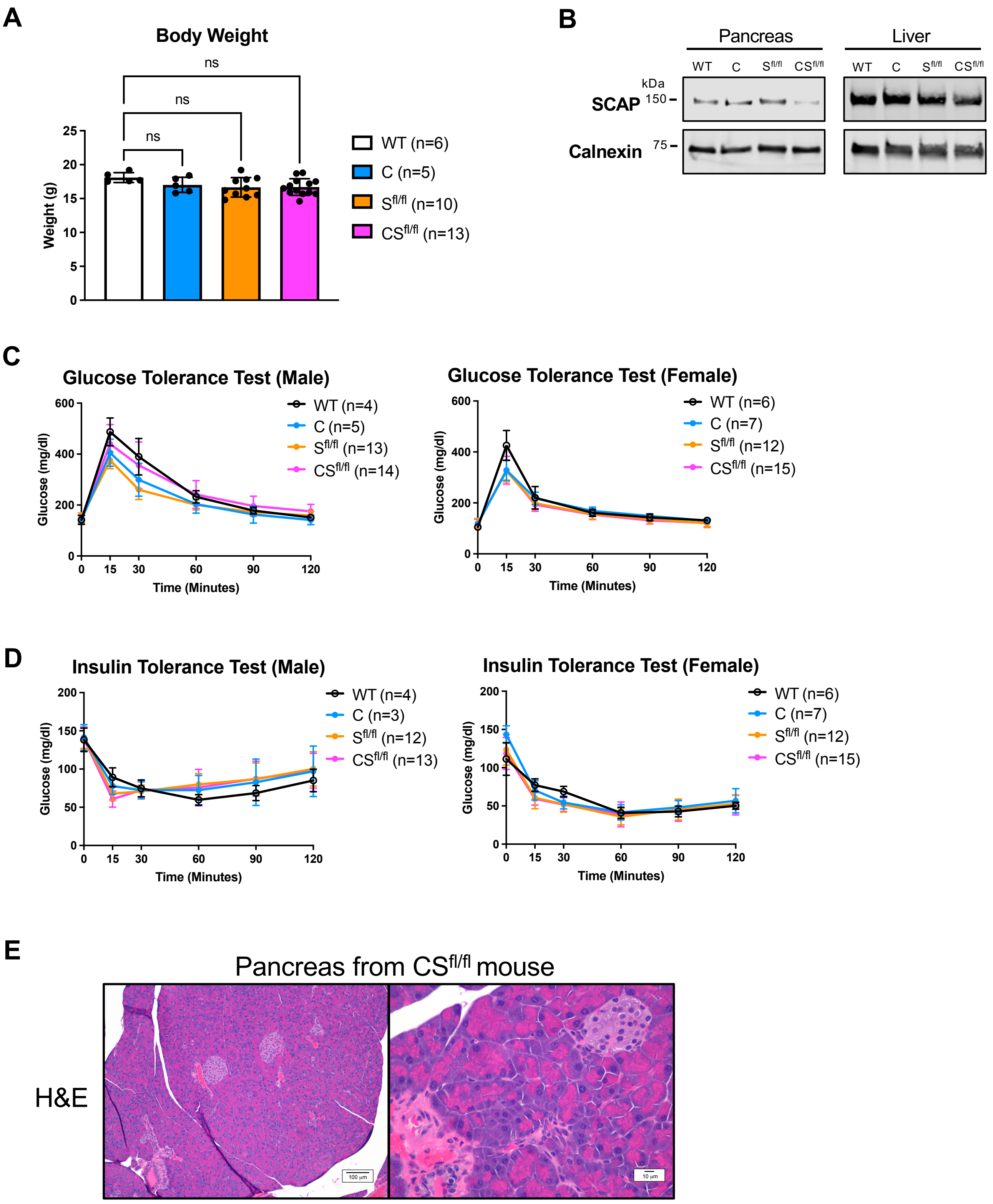

**FIGURE S1 – *Scap* is not required for development or function of the mouse pancreas.**

**A)** Body weights of 6-week-old female mice of C57BL/6 wildtype (WT), Pdx1-Cre (C), $Scap^{fl/fl}$ ($S^{fl/fl}$), or Pdx1-Cre $Scap^{fl/fl}$ ($CS^{fl/fl}$) mice. Statistical significance was determined by one-way ANOVA. Error bars denote standard deviation. **B)** Immunoblot analysis of microsomal membranes (40 µg) isolated from mouse pancreas and liver from indicated genotypes (1 mouse per sample). Blots were probed for SCAP and calnexin served as a loading control. **C)** Glucose tolerance tests in 6-week-old male (left graph) and female (right graph) mice with indicated genotypes. Error bar denotes standard deviation. **D**) Insulin tolerance test in 6-week-old male (left graph) and female (right graph) mice with indicated genotypes. Error bar denotes standard deviation. **E)** Representative H&E sections of the pancreas from a $CS^{fl/fl}$ mouse at low magnification (10X, left image) and higher magnification (40X, right image).

### Supplementary Figure 2

**A**

| Mouse # | Genotype |
|---|---|
| 1 | C |
| 2 | $CS^{fl/+}$ |
| 3 | $CS^{fl/fl}$ |
| 4 | KPC |
| 5 | $KPCS^{fl/+}$ |
| 6 | $KPCS^{fl/fl}$ |

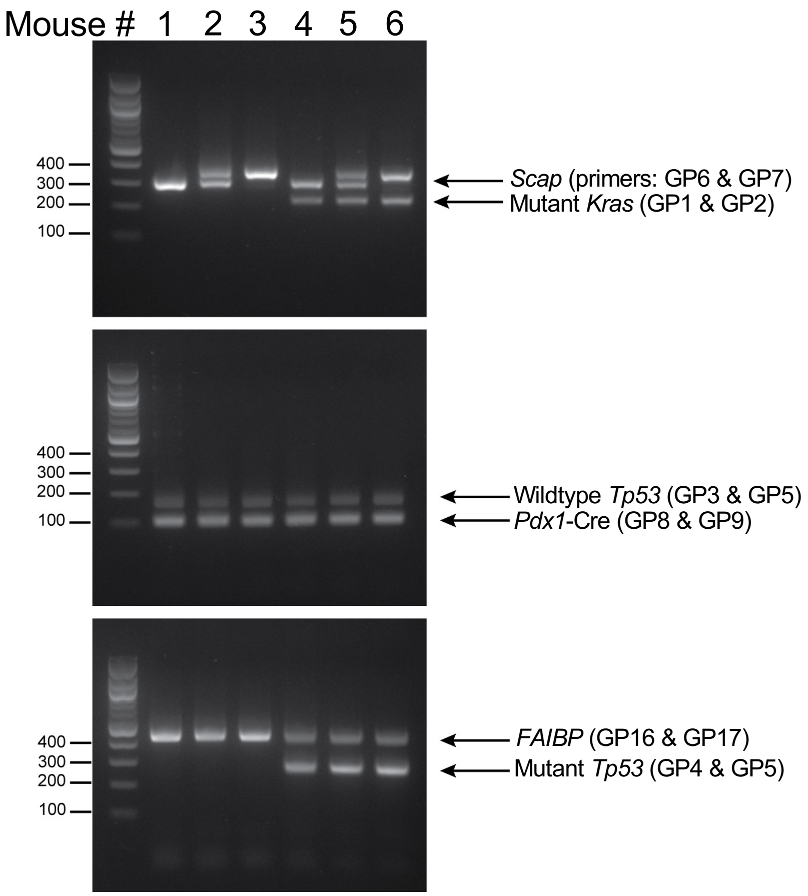


**B**

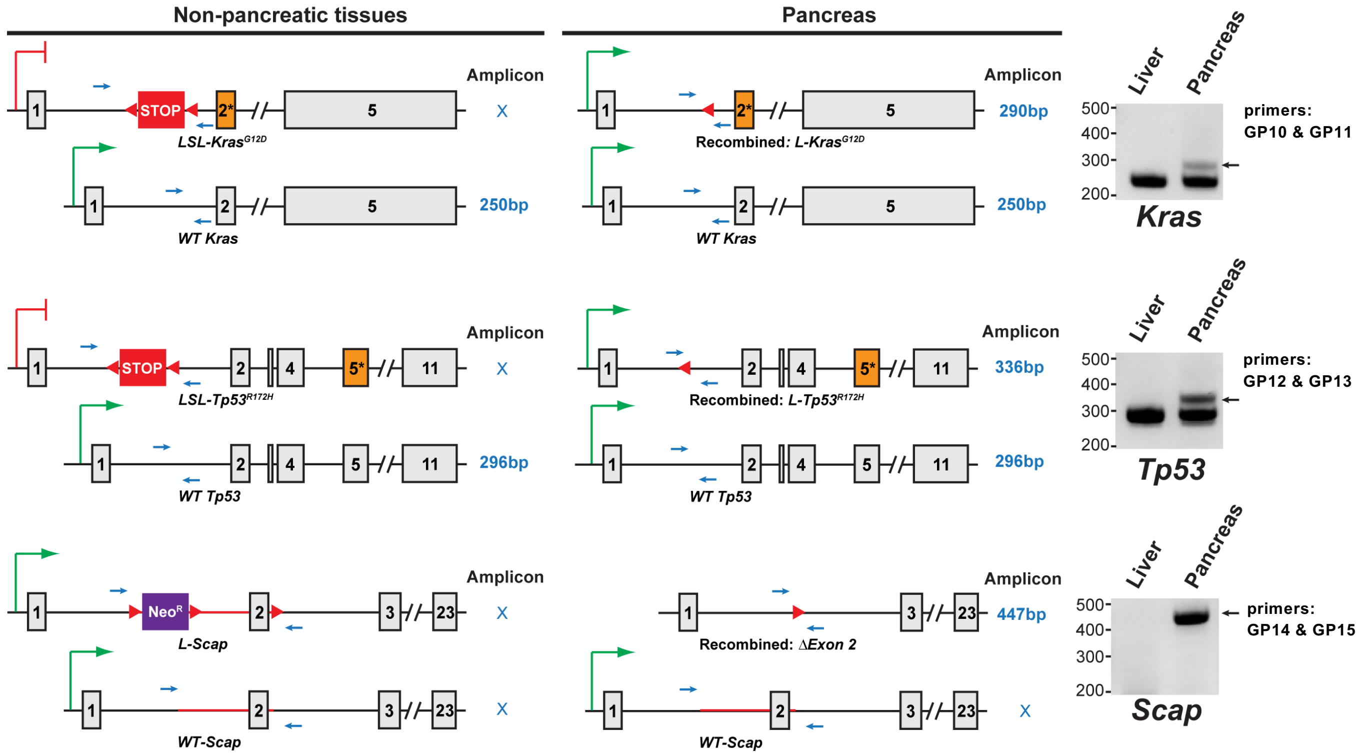

### Supplementary Figure 2 continued

**C**

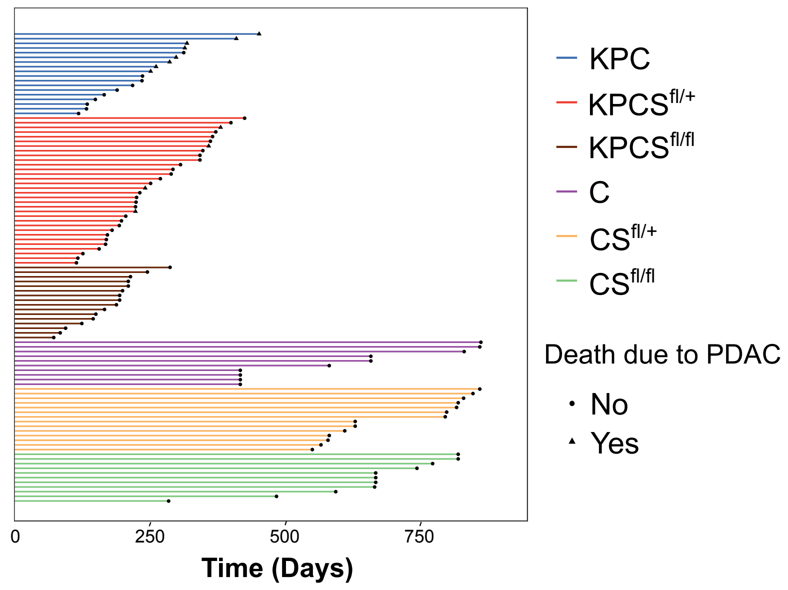


**D**

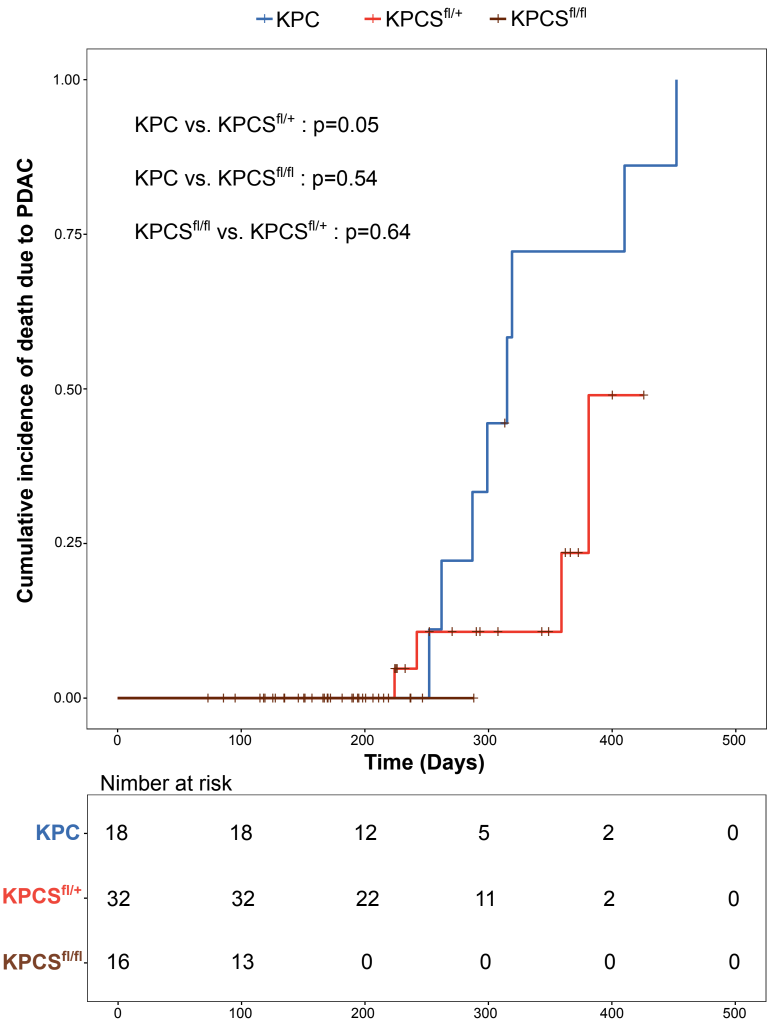

**FIGURE S2 – Mouse genotyping methods and survival study data.**

**A)** Representative genotyping of non-pancreatic tissue (tail or ear punch) of all mouse cohorts for identification purposes. *FAIBP* is fatty acid intestinal binding protein, an internal control. **B**) Representative genotyping of pancreas and liver tissue to identify specific genetic recombination events in *Kras*, *Trp53*, and *Scap*. PCR bands were not observed when the amplicons were too large to be amplified using the defined extension time (25 seconds). In the schematic of *Scap* recombination, exon numbers were modified based on updated NCBI RefSeq database. It was originally described that *loxP* sites are located upstream of exon 1 and in intron 1 (24). **C)** Swimmer plot showing lifespan of subjects in study and whether death was due to PDAC. Lifespan in days was calculated using date of birth and date of death. **D)** Figure shows the cumulative incidence curves of death due to PDAC for each genotype group. For this analysis, we considered death due to other causes as a censoring event. Log rank test was used for pairwise comparisons.

### Supplementary Figure 3

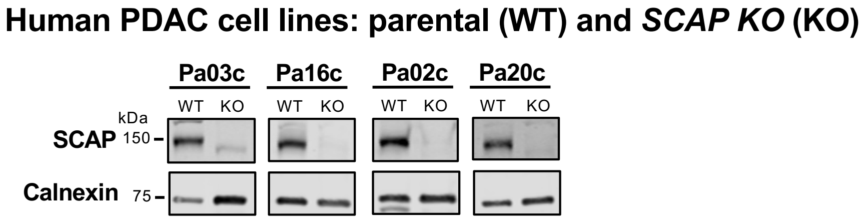

**FIGURE S3 – Generation of human PDAC *SCAP* knockout cell lines.**

Immunoblot analysis of Pa02c, Pa03c, Pa16c, and Pa20c cells for SCAP. Wildtype (WT) and *SCAP* knockout cells (KO) were cultured in 10% FBS supplemented medium. Membrane-enriched extracts (50 µg) were harvested and probed for SCAP. Calnexin served as a loading control.

#### Supplementary Figure 4

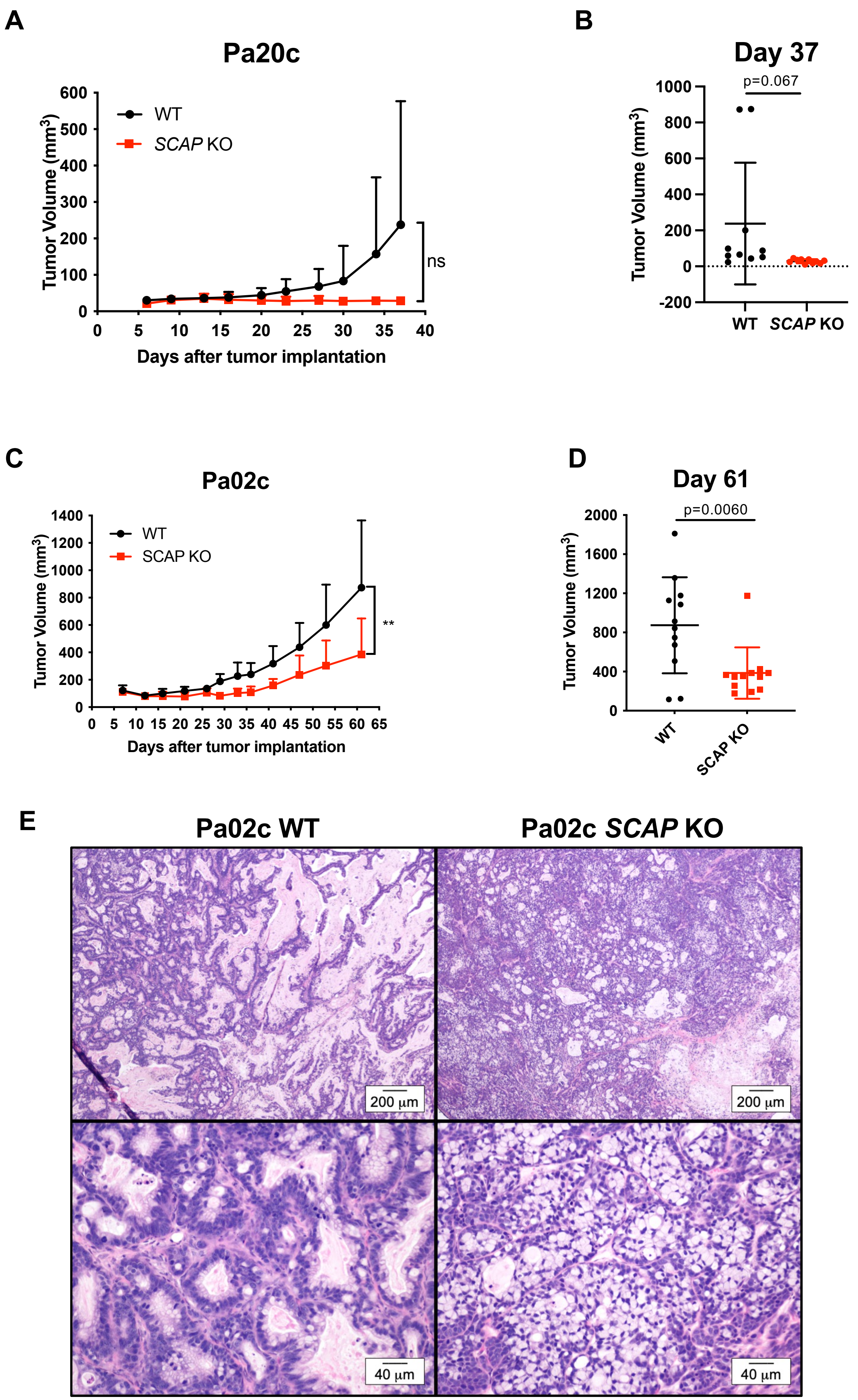

**FIGURE S4 – *SCAP* is required for human PDAC tumor growth in mouse orthotopic xenograft models.**

**A)** Nude mice were subcutaneously injected with $1x10^6$ Pa20c cells in both flanks (two tumors per mouse). Once visible, tumors were measured, and volume calculated. Each group contained 5 mice. Data represent mean ± SD (ns, not significant, student's t-test). **B)** Individual tumor volumes at day 37, n=10 tumors per group. **C)** Nude mice were subcutaneously injected with $1x10^6$ Pa02c cells in both flanks (two tumors per mouse). Once visible, tumors were measured, and volume calculated. Each group contained 6 mice. Error bar denotes standard deviation. Statistical significance was determined using student's t-test. (**, $p<0.01$). **D)** Individual tumor volumes at day 61. Error bar denotes standard deviation. Statistical significance was determined using student's t-test. **E)** Representative hematoxylin and eosin (H&E) stained sections of formalin fixed tumor tissues from the mice in **A** and **C** showing a low magnification (4X) image (upper row) and a high magnification (20X) image (bottom row) of tumor sections.

### Supplementary Figure 5

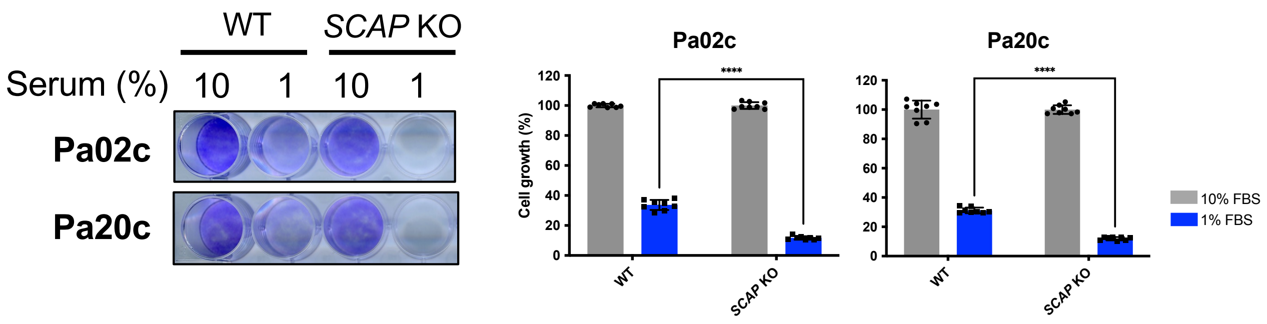

**FIGURE S5 – *SCAP* is required for PDAC cell growth and survival in low serum conditions.**

Cell growth assay of Pa02c and Pa20c cell lines. WT and *SCAP* knockout cells were cultured in either 10% FBS or 1% FBS for 7 days. Plates were stained with crystal violet as a measure of cell proliferation. Quantification of crystal violet staining is shown (n = 3 per group). For each cell line, growth was normalized to the 10% FBS condition. Statistical significance was determined using two-way ANOVA and Tukey's test. P values are indicated: < 0.05 (*); < 0.01 (**); < 0.001 (***); < 0.0001 (****), not significant (ns). Error bar denotes standard deviation.

### Supplementary Figure 6

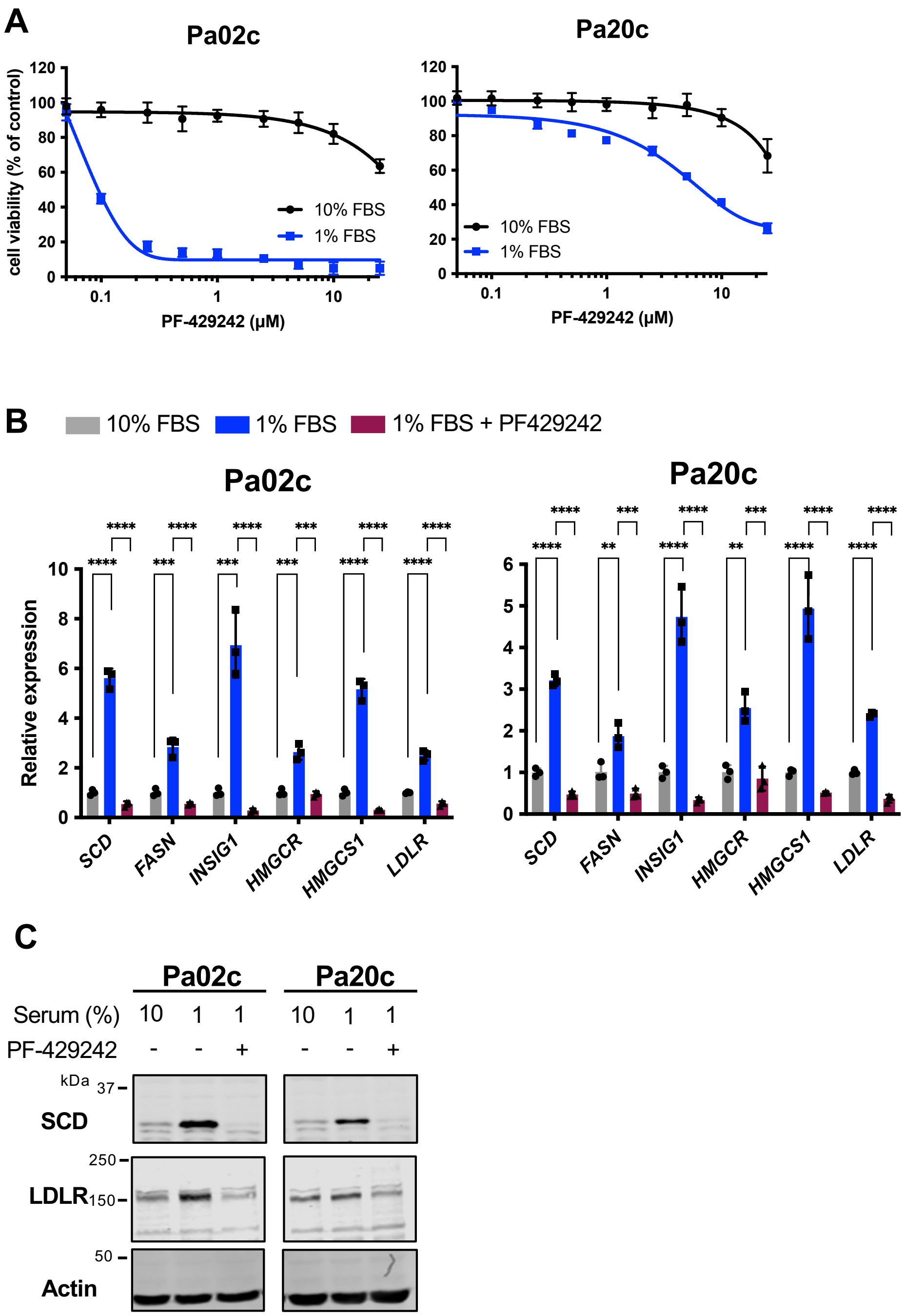

**FIGURE S6 – Site-1 protease activation of the SREBP pathway is required for PDAC cell growth in low serum conditions.**

**A)** PDAC cells (Pa02c and Pa20c) were cultured in either 10% or 1% FBS with indicated concentrations of the Site-1 protease inhibitor PF-429242 for 72 hours, and cell growth was determined using a MTS assay with data normalized to 10% FBS untreated cells. Data are representative of 2 biological replicates with 3 technical replicates for each biological replicate. Error bar denotes standard deviation. **B)** Pa02c and Pa20c cells were cultured in 10% FBS, 1% FBS, or 1% FBS with PF-429242 (10 mM) for 16 hours. Quantitative real-time PCR was performed for SREBP-1 (*SCD, FASN, INSIG1*) and SREBP-2 (*HMGCR, HMGCS1, LDLR*) target genes. *GAPDH* served as a control. Data are representative of 2 biological replicates with 3 technical replicates for each biological replicate. Error bar denotes standard deviation. Statistical significance was determined using one-way ANOVA and Tukey's HSD test. P values are indicated: < 0.05 (*); < 0.01 (**); < 0.001 (***); < 0.0001 (****), not significant (ns). **C)** Immunoblot analysis of whole cell lysates from Pa02c and Pa20c cells for SREBP target protein expression. Cells were cultured under the same conditions as in B. Whole cell protein extracts were harvested and probed for either SCD or LDLR. Actin served as a loading control. The figure is representative of 2 biological replicates.

### Supplementary Figure 7

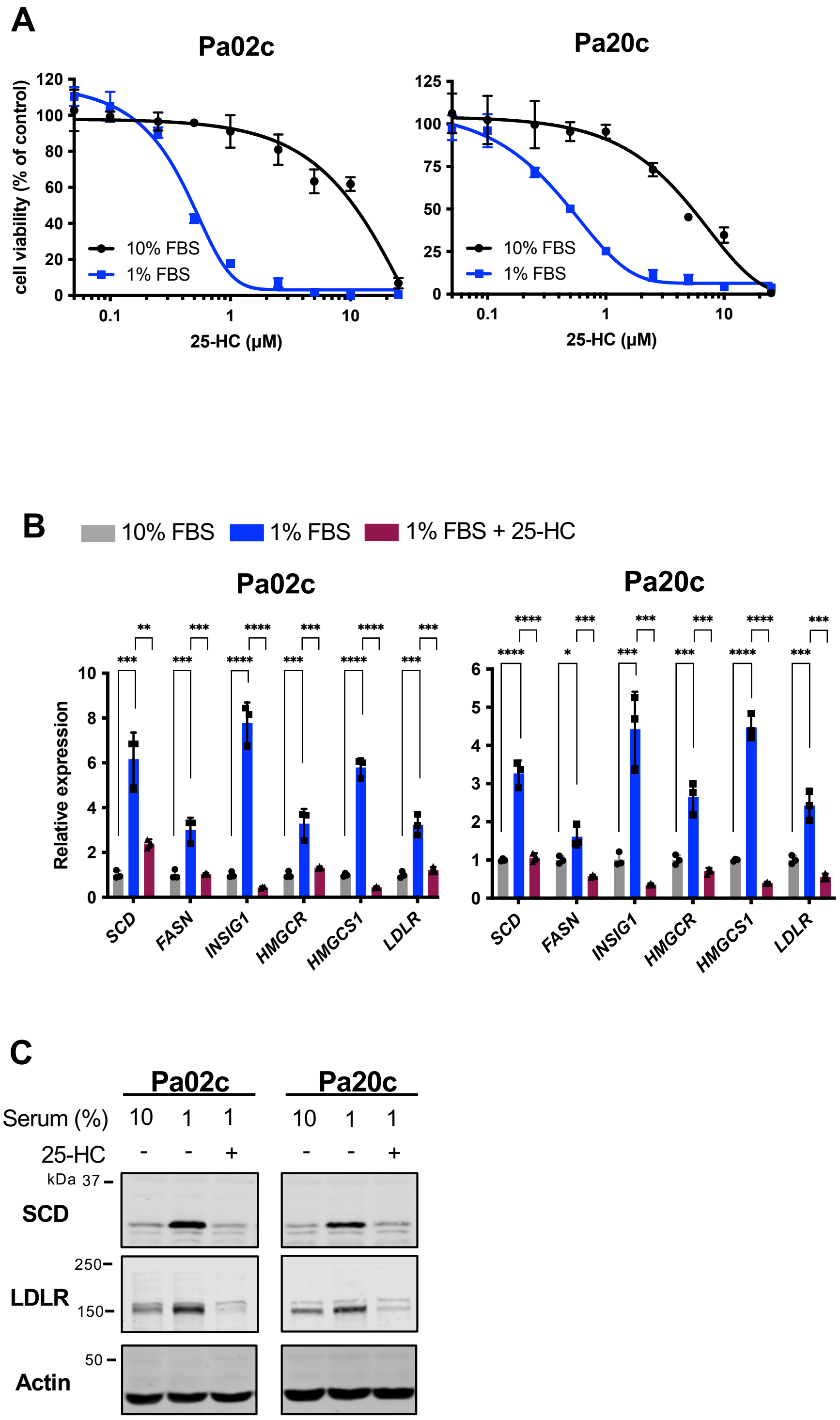

**FIGURE S7 – SREBP pathway activation is required for PDAC cell growth in low serum conditions.**

**A)** PDAC cells (Pa02c and Pa20c) were cultured in either 10% or 1% FBS with indicated concentrations of 25-hydroxycholesterol (25-HC) for 72 hours, and cell growth was determined using MTS assay and normalized to 10% FBS untreated cells. Data are representative of 2 biological replicates with 3 technical replicates for each biological replicate. **B)** Pa02c and Pa20c cells were cultured in either 10% FBS, 1% FBS, or 1% FBS containing 25-hydroxycholesterol (2.5 μM) for 16 hours. Quantitative real-time PCR was performed for target genes of SREBP1 (*SCD, FASN, INSIG1*) and SREBP2 (*HMGCR, HMGCS1, LDLR*). *GAPDH* served as a control. Data are from 2 biological replicates with 3 technical replicates for each biological replicate. Error bar denotes standard deviation. Statistical significance was determined using one-way ANOVA and Tukey's HSD test. P values are indicated: < 0.05 (*); < 0.01 (**); < 0.001 (***); < 0.0001 (****), not significant (ns). **C)** Immunoblot analysis of Pa02c and Pa20c whole cell lysates for indicated SREBP target protein expression. Cells were cultured under the same conditions as in B. Protein extracts were harvested and probed for either SCD or LDLR. Actin served as a loading control. The figure is representative of 2 biological replicates.
